# Gestational LSD exposure in mouse rapidly reaches embryonic CSF and is associated with altered choroid plexus signaling, cerebral cortical development, and offspring behavior

**DOI:** 10.1101/2025.09.30.677638

**Authors:** Ya’el Courtney, Josephine M. Anderson, Christian Lagares-Linares, Cody J. Wenthur, Maria K. Lehtinen

**Affiliations:** Department of Pathology, Boston Children’s Hospital and Harvard Medical School; Boston, MA, USA; Program in Neuroscience, Harvard Medical School; Boston, MA, USA; School of Pharmacy, University of Wisconsin; Madison, WI, USA; Transdisciplinary Center for Research in Psychoactive Substances, University of Wisconsin; Madison, WI, USA

## Abstract

Classic serotonergic psychedelics engage 5-HT receptors throughout the nervous system, but how maternal exposure intersects with embryonic brain interfaces is poorly defined. Here we tested in mice whether maternally administered lysergic acid diethylamide (LSD) accesses embryonic cerebrospinal fluid (CSF) and whether embryonic choroid plexus (ChP) – a CSF-secreting epithelium enriched for *Htr2c* – mounts an acute response. Following a single maternal injection (0.3 mg kg⁻¹, subcutaneous), LSD was detectable in embryonic CSF within 5–15 minutes at E12.5 and E16.5. Thirty minutes after maternal dosing, LSD induced Fos in embryonic ChP across ventricles and was accompanied by rapid apical remodeling and increased embryonic CSF protein. In parallel cohorts, psilocybin, 5-MeO-DMT, and the 5-HT₂C agonist WAY-161503 elicited a similar Fos response in ChP. Prenatal LSD exposure during mid-gestation was associated with altered S1 cortical cellularity and projection-neuron subtype marker composition at P8; regimen-dependent effects included male-biased changes in SATB2⁺ and CTIP2⁺ populations after repeated exposure. In adulthood, offspring exhibited modest, male-predominant reductions in prepulse inhibition and increased rotational stereotypy. Together, these data identify embryonic CSF as a rapidly accessible compartment for maternal LSD and support a model in which serotonergic agonists can acutely engage ChP epithelium during cerebral cortical development.

**Significance:** Psychedelic use during pregnancy is increasing, but the speed and extent to which these drugs access the embryonic CNS remain unknown. We show that a single maternal dose of LSD appears in mouse embryonic cerebrospinal fluid within five minutes and provokes an immediate response in the choroid plexus, a serotonin receptor-rich epithelium that regulates CSF composition. Psilocybin, 5-MeO-DMT, and a selective 5-HT₂C agonist trigger a similar response. Mid-gestational exposure alters cortical neuron composition in neonates and produces persistent behavioral abnormalities in adult offspring, including stereotypies evident from weaning. These data reveal that maternal serotonergic agonists rapidly access embryonic CSF, acutely activate choroid plexus epithelium, and are associated with lasting changes in cortical composition and offspring behavior.

## Main Text

Serotonin shapes neural proliferation, migration, and circuit assembly across embryonic and early postnatal development (Gaspar et al., 2003; Bonnin et al., 2011; Shah et al., 2018; Carvajal-Oliveros and Campusano, 2021). Exogenous serotonergic agonists administered during gestation therefore have the potential to engage developmental programs, but the routes by which maternal compounds access embryonic brain compartments - and which embryonic interfaces respond acutely - remain incompletely defined. A key unresolved question is whether maternally administered serotonergic psychedelics reach embryonic cerebrospinal fluid (CSF), where they could interact with CSF-contacting target cells during cerebral cortical neurogenesis.

One candidate target interface is the choroid plexus (ChP). The ChP is a secretory epithelium that produces CSF (Damkier et al., 2013; Lun et al., 2015; MacAulay et al., 2022) and helps shape the molecular environment of the developing brain (Saunders et al., 2023). Transcriptomic atlases indicate that *Htr2c* is prominently expressed in ChP epithelium relative to other serotonin receptor subtypes (Dani et al., 2021), and early autoradiographic work showed that ^125^I-LSD binds serotonergic sites on ChP epithelial cells at a density ten-fold higher than any other serotonergic site in brain homogenates (Yagaloff and Hartig, 1985). Functionally, cultured ChP epithelial cells mount immediate-early gene responses to serotonergic hallucinogens (Sanders-Bush and Breeding, 1991). In parallel, CSF-borne cues influence cerebral cortical development (Gato and Desmond, 2009; Lehtinen et al., 2011; Fame and Lehtinen, 2020), raising the possibility that acute drug-evoked changes in ChP state could modulate developmental signaling via CSF.

We focused on LSD as a prototypical, high-potency serotonergic psychedelic with well-characterized receptor pharmacology and extensive rodent behavioral and pharmacokinetic data, which facilitated sensitive quantification from the small volumes of embryonic CSF. To test whether ChP activation is specific to LSD versus a broader feature of serotonergic agonism, we additionally compared acute ChP Fos responses following maternal exposure to psilocybin, 5-MeO-DMT, and the selective 5-HT₂C agonist WAY-161503.

Here we asked three questions. First, does maternally administered LSD rapidly access embryonic CSF during mid-gestation? Second, does the embryonic ChP mount an acute transcriptional and structural response consistent with serotonergic receptor engagement? Third, are mid-gestational exposure regimens associated with changes in early postnatal cerebral cortical cellular organization and adult behavioral readouts in offspring? We combine LC–MS/MS pharmacokinetics with embryonic CSF sampling, ChP immediate-early gene and epithelial remodeling assays, and analyses of cortical marker composition and behavior. Our data support an interface-focused model for how maternal serotonergic agonists can engage the developing brain.

## Materials and Methods

See Supplemental Methods for a complete description of experimental methods as well as detailed information about all resources used and material associated with this article.

### Ethical approval

All procedures complied with institutional and national guidelines. Animal work was approved by the Boston Children’s Hospital IACUC.

### Animals

Adult and timed-pregnant CD-1 mice (Charles River; Jackson Laboratory; or in-house colonies) were maintained at 21 ± 1.5 °C, 35–70% humidity, 12-h light/dark with ad libitum food/water. Unless stated, experiments used equal numbers of males and females. Sample sizes, sex, and ages are provided in figure legends. Litter was tracked and used as a factor in sensitivity analyses.

### Drugs and dosing (in vivo)

Timed-pregnant dams received subcutaneous injections in sterile 0.9% saline: WAY-161503 3 mg kg⁻¹ (Tocris, Cat. 1801); (+)-lysergic acid diethylamide tartrate (LSD) 0.3 mg kg⁻¹ (NIDA Drug Supply Program); psilocybin 3 mg kg⁻¹ (NIDA); 5-MeO-DMT 50 mg kg⁻¹ (NIDA). Solutions: WAY-161503, 10 mM at 1 µL g⁻¹; LSD tartrate, 0.6 mM at 0.6 µL g⁻¹; psilocybin from a 15.03 mg mL⁻¹ stock at 0.1995 µL g⁻¹ (saline added to a minimum 50 µL total); 5-MeO-DMT, 50 mg mL⁻¹ at 1 µL g⁻¹. Body weight and injection/sacrifice times were logged for PK.

### Choroid plexus (ChP) dissections

Lateral ventricle (LV), third ventricle (3V), and fourth ventricle (4V) ChP were microdissected using standard landmarks. LV ChP was exposed by hippocampal reflection; adult 3V ChP was visualized along the dorsal midline; 4V ChP was accessed via cisterna magna exposure.

### Behavioral testing

Behavior was performed with the BCH Animal Behavior & Physiology Core (blinded, randomized; litter-balanced where possible). Open-field test: Adult CD-1 mice (n=10 per condition unless noted) explored a 40×40 cm arena for 15 min. Primary outcomes were total distance traveled and rotational stereotypy; spatial occupancy metrics (e.g., centroid/heatmaps) were used as descriptive visualizations. Rotations were quantified from the animal’s instantaneous heading angle relative to the arena center. A “rotation” was defined as ≥360° cumulative angular displacement in a single direction, with direction determined by the sign of angular velocity. Prepulse inhibition: Background 60 dB, pre-pulses 70/74/78/82 dB (20 ms), startle 105 dB (40 ms), 100 ms ISI, 10–30 s randomized ITI. %PPI = 100 × [(pulse-alone – (prepulse+pulse))/pulse-alone].

### Immunostaining and imaging

Explants: ChP explants fixed (4% PFA, 10 min), permeabilized (0.1% Triton X-100), primary overnight at 4 °C, secondary 2 h at room temperature; Hoechst 33342 counterstain; Fluoromount-G mounting. Sections: Postnatal brains perfused with PBS then 4% PFA, cryoprotected (10%→20%→30% sucrose), embedded in OCT, permeabilized, and stained as above. Confocal imaging on Zeiss LSM710/980 (20× air; 63× oil). Acquisition in ZEN Black; Airyscan processing in ZEN Blue when specified.

### Image quantification

Aposomes: Four 20× fields per explant were scored by a blinded rater; the mean per explant constituted N=1. FOS+ nuclei: Four 20× fields per explant were thresholded in FIJI; Analyze Particles counted FOS and DAPI objects (>200 px). Proportion FOS+ = FOS/DAPI; field means per explant constituted N=1. Cortical marker classes: SATB2/CTIP2/TBR1 images spanning S1 cortex (L1 to corpus callosum) were exported as 4-channel TIFFs and analyzed in Biodock (trained model; QC-checked). Cells were binned into six equal depth bins (y-axis). Class frequencies per bin were exported to R for summaries and to GraphPad Prism for plotting.

### Real-time quantitative PCR (RT-qPCR)

ChP was microdissected, snap-frozen, and total RNA extracted (NEB Monarch Total RNA kit). 100 ng RNA was reverse-transcribed (NEB LunaScript). qPCR used TaqMan Gene Expression Master Mix (Thermo Fisher) with Mm00487425_m1 (Fos) and Hs99999901_s1 (18S) on a Roche LightCycler 480II. Cycling: 95 °C 10 min; 40×(95 °C 15 s/60 °C 1 min). Triplicates were run. Relative expression used 2^–ΔΔCT (normalized to 18S).

### Whole-mount clearing and light-sheet imaging

E12.5 heads were fixed in 4% PFA, processed with SHIELD (LifeCanvas; small-sample workflow), delipidated in SDS buffer at 45 °C, immunolabeled for c-FOS, index-matched in EasyIndex, and imaged on a Zeiss Lightsheet 7 (5× objective). Z-stacks were stitched in ZEN; 16-bit TIFFs were volume-rendered in syGlass v2.3.0 with identical linear transfer functions.

### Pharmacokinetics Analysis via LC–MS/MS

Serum and CSF samples were analyzed using a Sciex QTrap 5500 coupled to a Waters ACQUITY UPLC I-Class. LSD and d3-LSD (ISTD) were quantified using quadratic standard curves, 1/x^2 weighting; calibrators retained if ≥85% (R²≥0.995). Mobile phases were water (A) and acetonitrile (B) with 0.1% Formic Acid, run a 0.375mls/min through a Phenomenex Kinetex Phenyl-Hexyl (1.7um, 100x1.2mm) held at 35 °C. Gradient: 5–40% B to 1.75 min; 95% B to 2.5 min; re-equilibrate to 3.0 min. Analytes were monitored in MRM mode (positive) via the following unique transitions; (324.162>223.00, 209.10, 207.10) and d3-LSD (327.160> 226.10, 208.00, 180.10). Calibrators were generated using blank CD1 mouse serum and artificial CSF (aCSF). Samples were prepared via a Waters Sirocco Protein Precipitation Plates. Injections were submitted in triplicate in randomized sets of 6 with a blank between each injection.

### Quantification and statistical analysis

Analyses were pre-planned, randomized, and blinded where possible. Power analyses (α=0.05, 1–β=0.80) informed group sizes using pilot variance. Outliers: ROUT (Q=1%). Normality (Shapiro–Wilk) and variance homogeneity (F-test/Bartlett) guided test selection. Parametric: unpaired t-test (Welch if unequal variances); one-way ANOVA (Tukey) for ≥3 groups; two-way ANOVA (Sidak) for two factors. Nonparametric: Mann–Whitney U; Kruskal–Wallis (Dunn). Sex was included where relevant; litter evaluated as a random intercept in sensitivity models. Data are mean ± SD (or mean ± SEM for multiple measures per animal). Exact n, tests, and P values are reported in legends.

## Results

### Maternal LSD rapidly reaches embryonic CSF in mouse

We tested and found that maternally administered LSD crosses the placenta and reaches the embryo by measuring drug levels in maternal serum, embryonic serum (E16.5), and embryonic CSF at E12.5 and E16.5 following a single subcutaneous dose of 0.3 mg kg⁻¹; samples were collected at 5, 15, 30, 60, and 120 min (**Fig. 1A**). Doses were selected to ensure target engagement in mouse and are not human-equivalent. Maternal serum was obtained via cardiac puncture. At E16.5, embryonic serum was collected by venipuncture of a cervical vein under stereomicroscopic guidance using a pulled glass microcapillary pipette. Embryonic CSF was collected at E12.5 and E16.5 from the cisterna magna using a glass microcapillary pipette under stereomicroscopic guidance, as previously described (Zappaterra et al., 2013), minimizing blood or neuroepithelial contamination.

**Fig. 1.**
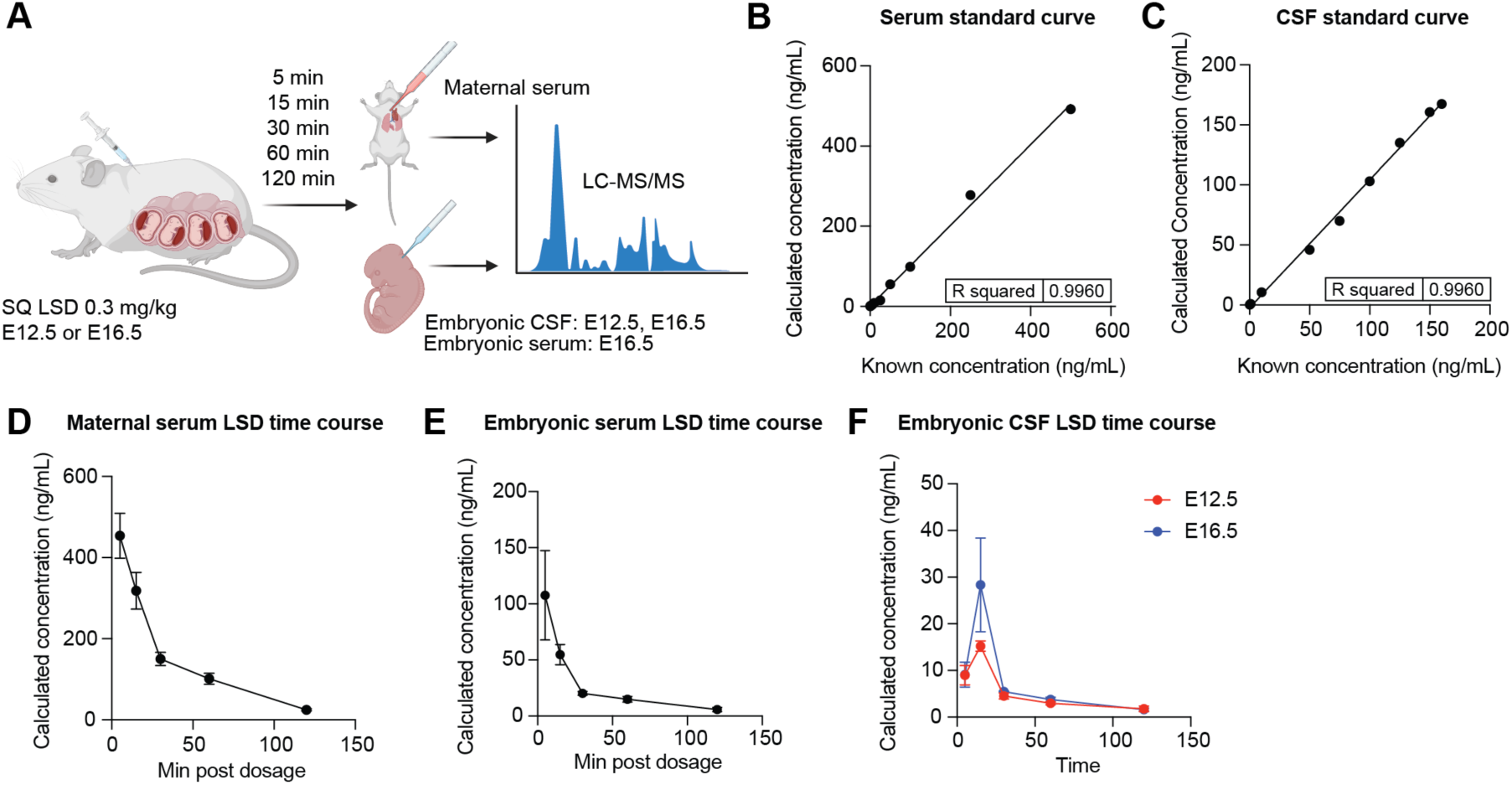
Maternal LSD rapidly enters embryonic circulation and CSF. **(A)** Schematic of the pharmacokinetic study. Pregnant mice received a single subcutaneous injection of LSD (0.3 mg kg⁻¹) at E12 .5 or E16.5. Maternal serum, embryonic serum (E16.5) and embryonic CSF (E12.5, E16.5) were sampled at 5, 15, 30, 60, and 120 min and analyzed by LC-MS/MS. **(B)** Calibration curve for LSD spiked into serum (R² = 0.996). **(C)** Calibration curve for LSD spiked into CSF (R² = 0.996). **(D)** Time course of LSD concentration in maternal serum after 0.3 mg kg⁻¹ LSD (mean ± SEM, n = 3 dams per time point). **(E)** Time course of LSD concentration in E16.5 serum after 0.3 mg kg⁻¹ maternal LSD (mean ± SEM, n = 3 pooled litters per time point). **(F)** LSD concentration in embryonic CSF at E12 .5 or E16.5 peaks at 15 min then declines yet remains detectable at 120 min (mean ± SEM, n = 3 pooled litters per time point).

We developed a small-volume LC–MS/MS assay for mouse serum and embryonic CSF using d3-LSD as the internal standard. Calibration curves were prepared in-matrix (blank serum or artificial CSF), were linear across the working range (R² ≈0.997), and supported a lower limit of quantitation of 0.25 ng mL⁻¹. Calibration curves prepared in-matrix met acceptance criteria, with back-calculated bias within −8% to +9% (serum) and −8% to +8% (CSF) and replicate CV ≤ 2.7% both matrices (**Fig. 1B–C**).

Following a single maternal dose of 0.3 mg kg⁻¹, maternal serum reached its highest concentration at 5 min (419.6 ng mL⁻¹) and then declined with an elimination half-life of ≈39.0 min based on the 30–120 min log-linear phase (**Fig. 1D**). At E16.5, embryonic serum peaked at 5 min (≈107.8 ng mL⁻¹; ∼26% of the maternal maximum) and showed a half-life of ≈53.0 min by the same approach (**Fig. 1E**). Embryonic CSF contained measurable LSD by 5–15 min at both ages, with 15-min peaks of ≈15.2 ng mL⁻¹ (E12.5; t₁/₂ ≈64.9 min) and ≈28.4 ng mL⁻¹ (E16.5; t₁/₂ ≈52.5 min), remaining above the assay limit at 120 min (≈1.8 and 1.6 ng mL⁻¹, respectively) (**Fig. 1F**). At 15 min, CSF levels at E16.5 were ∼1.9× those at E12.5 and were ∼50% of the paired embryonic-serum concentration. Across 0–120 min, total embryonic-serum exposure was ∼17% of maternal serum, and CSF exposure at E16.5 exceeded E12.5 by ∼1.4×. Taken together, these data demonstrate that LSD crosses the placenta and reaches the embryonic CSF within minutes.

### Prenatal LSD reshapes S1 neuron composition and microglia at P8

Maternal LSD exposure was associated with regimen-dependent changes in S1 cortical cellularity and projection-neuron subtype marker composition at P8 (**Fig. 2**). The experimental design (**Fig. 2A**) contrasted a single 0.3 mg kg⁻¹ injection at E12.5 (“acute”) with the same dose given once daily from E12.5-E16.5 (“repeated”). At the 0.3 mg kg⁻¹ dose, no adverse events such as maternal weight loss, altered grooming, or changes in pup retrieval were observed. Litter size, pup viability, birth weight, brain weight, and brain-to-body ratio at birth did not differ between LSD-exposed and control litters (**Fig. S1A-D**). Brains were examined at postnatal day 8 (P8), when laminar identities and glial profiles are well resolved but before extensive synaptic pruning.

**Fig. 2.**
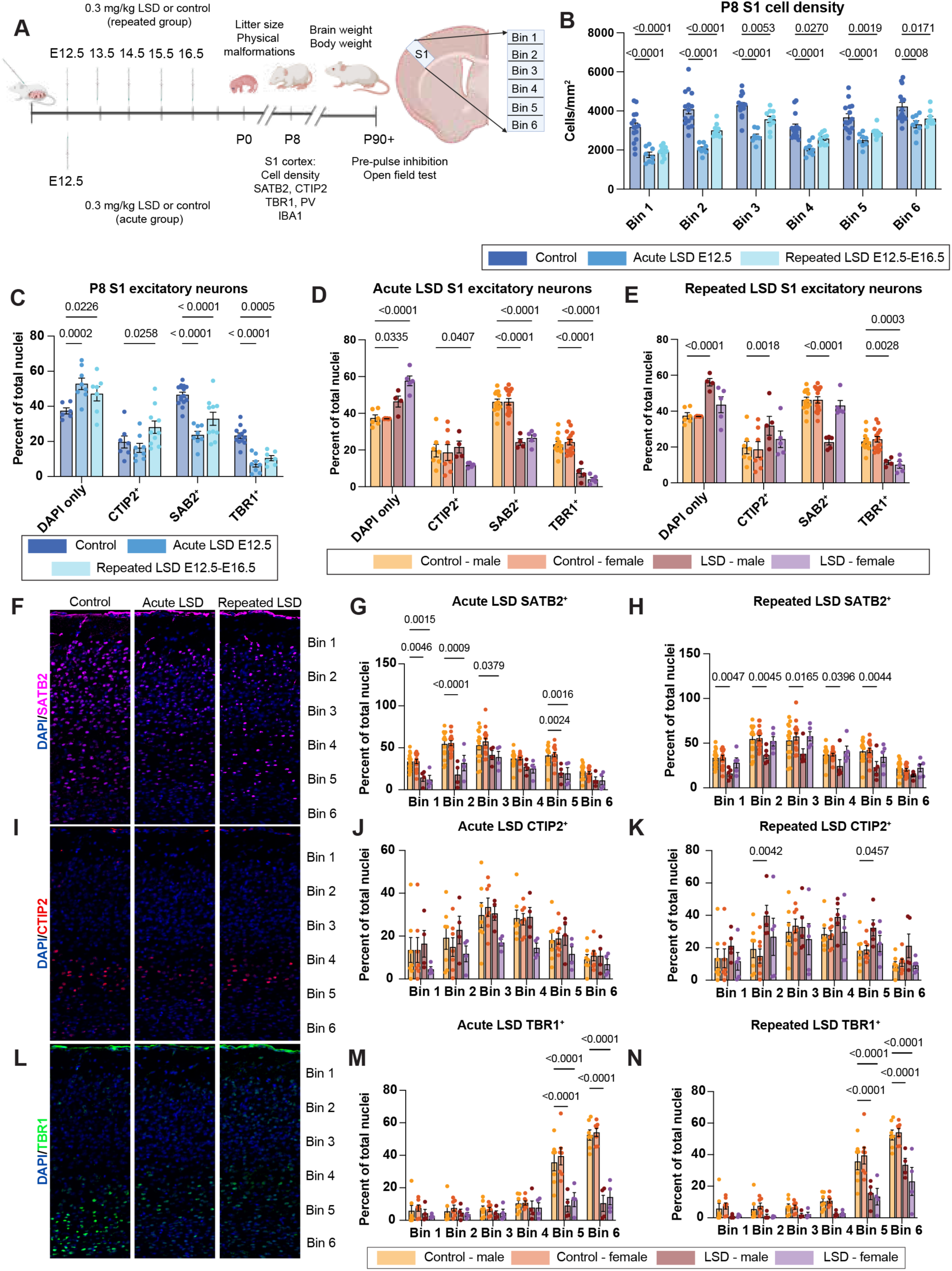
Prenatal LSD alters S1 cortical composition with regimen-dependent and partly sex-dependent effects. **(A)** Experimental design and S1 sampling. Pregnant dams received a single LSD dose at E12.5 (“acute”) or daily doses at E12.5–E16.5 (“repeated”) at 0.3 mg kg⁻¹. Brains were analyzed at P8. S1 was divided into six equal-width bins from pia (Bin 1) to white matter (Bin 6). **(B)** Total cell density (DAPI⁺ nuclei mm⁻²) across depth. Both regimens reduce nuclei relative to controls; exact post hoc *P* values are printed above bins. **(C)** Sexes pooled: composition of glutamatergic projection-neuron subtype markers expressed as percent of total nuclei (DAPI-only, CTIP2⁺, SATB2⁺, TBR1⁺). Repeated exposure increases CTIP2⁺ and decreases SATB2⁺/TBR1⁺; acute exposure decreases SATB2⁺/TBR1⁺ without increasing CTIP2⁺ (panel *P* values shown). For panels **2B-C**, control n = 15 from 3 litters; Acute LSD n = 9 from 3 litters; Repeated LSD n = 10 from 3 litters. **(D–E)** Sex-stratified marker composition for acute (**D**) and repeated (**E**) exposure (color key indicates sex and treatment). **(F)** Representative S1 micrographs (male) stained for SATB2 (green) with DAPI (blue). Scale bar, 50 µm. **(G–H)** SATB2⁺ neurons by depth after acute (**G**) or repeated (**H**) exposure (percent of total nuclei per bin). Male-biased reductions are evident across superficial to mid layers; bin-wise *P* values are printed. **(I)** Representative S1 micrographs (male) stained for CTIP2 (green) with DAPI (blue). Scale bar, 50 µm. **(J–K)** CTIP2⁺ neurons by depth after acute (**J**) or repeated (**K**) exposure. Repeated exposure produces bin-restricted increases in males (bin-wise *P* values shown). **(L)** Representative S1 micrographs (male) stained for TBR1 (green) with DAPI (blue). Scale bar, 50 µm. **(M–N)** TBR1⁺ neurons by depth after acute (**M**) or repeated (**N**) exposure, showing deep-layer losses in both sexes (bin-wise *P* values shown). For panels 2**D-N**, control male n = 7 from 3 litters, control female n = 7 from 3 litters, acute LSD male = 4 from 3 litters, acute LSD female n = 5 from 3 litters, repeated LSD male n = 5 from 3 litters, repeated LSD female n = 5 from 3 litters.

Across six equal-width bins spanning pia to white matter, DAPI^+^ nuclei were reduced following LSD (two-way ANOVA, Treatment effect *P* < 0.0001; **Fig. 2B**). Acute exposure produced the larger deficit, 1,090-2,020 cells mm⁻² (21–49 % below control, post hoc *P* ≤ 0.0008 in every bin), whereas repeated exposure reduced nuclei by 580 – 1,260 cells mm⁻² (16–40 %; *P* ≤ 0.027). A Treatment × Sex interaction was not detected for total nuclei (F_2,168_ = 0.34, *P* = 0.71), indicating a comparable global loss in males and females.

SATB2, CTIP2, and TBR1 are transcription factors commonly used to operationalize major cortical projection-neuron subclasses during early postnatal development, and we used these markers as indices of subclass composition. Cerebral cortical layer marker quantification revealed shifts among corticofugal/subcerebral (CTIP2⁺), callosal (SATB2⁺) and corticothalamic (TBR1⁺) neurons (**Fig. 2C-E**). Repeated dosing increased the CTIP2⁺ fraction from 19.7 ± 1.5 % to 28.1 ± 1.9 % (difference +8.4 percentage points [pp], one-way ANOVA with Dunnett’s multiple comparisons *P* = 0.026), whereas acute dosing did not change CTIP2^+^ abundance. SATB2⁺ neurons were selectively vulnerable, dropping by 13.6 pp with repeated exposure (46.5 ± 1.6 % → 32.9 ± 2.1 %, *P* < 0.0001) and by 22.9 pp after acute exposure (to 23.7 ± 1.7 %, *P* < 0.0001). TBR1⁺ neurons declined under both regimens (–12.8 pp repeated, –16.1 pp acute; *P* ≤ 0.0005). The “DAPI-only” fraction (nuclei lacking these three markers) rose by 9.7 pp with repeated LSD (*P* = 0.023) and by 15.4 pp with acute LSD (*P* = 0.0002).

Stratifying by sex and cortical depth (**Fig. 2F–N**) revealed a significant Treatment × Sex × Bin interaction for CTIP2⁺ and SATB2⁺ populations (F_10,250_ = 3.21, *P* = 0.0007). In males, repeated LSD reduced SATB2⁺ neurons in bins 1–5 by 28–52 % (all *P* ≤ 0.040) and doubled CTIP2⁺ neurons in bin 2 (+20.5 pp, *P* = 0.0042), with smaller gains in bin 5. Females showed no detectable CTIP2^+^/SATB2^+^ shifts. TBR1⁺ loss was equivalent across sexes and confined to deep bins 5–6 (–19 – 22 pp; *P* < 0.0001). Acute exposure produced parallel SATB2⁺ and TBR1⁺ deficits in both sexes without altering CTIP2⁺ levels.

IBA1⁺ microglia density increased with repeated dosing from 166.8 ± 6.2 to 204.5 ± 8.3 cells mm⁻² (+22 %, Dunnett post hoc *P* = 0.0036, **Fig. S1E**) and rose modestly after acute dosing (192.6 ± 9.8 cells mm⁻², +16 %, *P* = 0.044, **Fig. S1F**). GFAP staining revealed reactive astrocytic rims surrounding ectopic parvalbumin-positive interneuron clusters posterior to the anterior commissure, which only appeared in LSD-exposed mice (**Fig. S1G**).

In sum, prenatal LSD exposure was associated with reduced S1 cellularity at P8 and altered composition of projection-neuron subtype marker populations. Both exposure regimens reduced SATB2⁺ and TBR1⁺ populations, while repeated exposure was additionally associated with increased CTIP2⁺ fractions in males and elevated microglial density. These data define regimen-and sex-contingent early postnatal cortical signatures following mid-gestational serotonergic exposure.

### Prenatal LSD exposure is associated with reduced sensorimotor gating (PPI) and increased rotational stereotypy

To test for lasting behavioral effects following in utero LSD exposure, we evaluated offspring at P90 on prepulse inhibition of acoustic startle (PPI, **Fig. 3A**) and open-field behavior. Pregnant mice received either one single LSD injection at E12.5 (acute) or daily injections from E12.5 to E16.5 (repeated), with repeated saline-treated dams as controls (**Fig. 3B**).

**Fig. 3.**
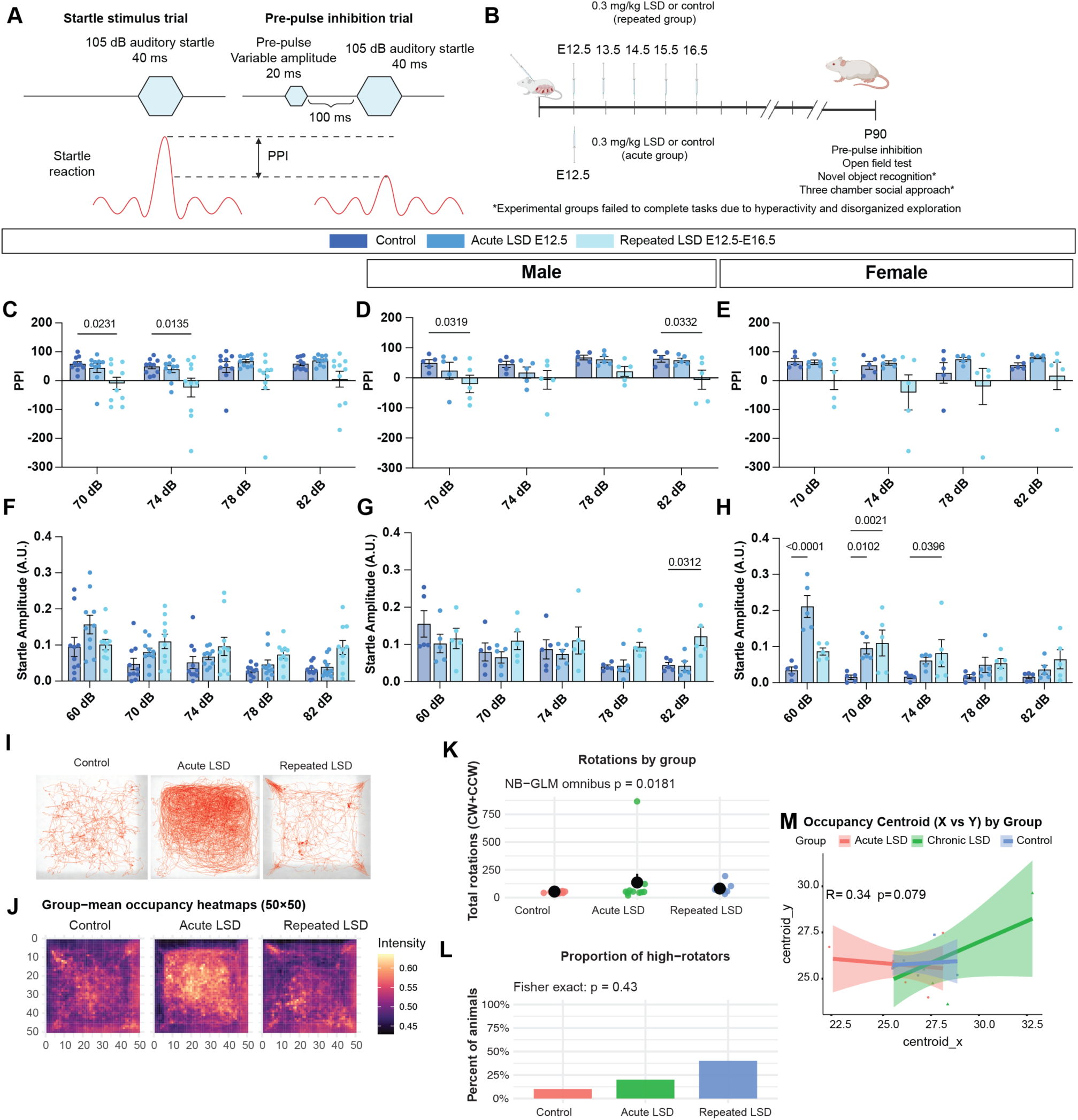
Embryonic LSD exposure reduces prepulse inhibition and increases rotational stereotypy. Analyses were blinded; litter was tracked and included as a random intercept in sensitivity models. For all behavioral assays presented, control n = 10 from 3 litters, acute LSD n = 10 from 3 litters, repeated LSD n = 10 from 3 litters; 5 male and 5 female mice per experimental condition. **(A)** PPI task schematic. **(B)** Maternal dosing and testing timeline (acute E12.5; repeated E12.5–E16.5; saline controls). **(C–E)** Prepulse inhibition (PPI) at P90: **(C)** sexes pooled, PPI reduced after prenatal LSD; **(D)** males, reduced PPI after repeated exposure; **(E)** females, no group-level difference at this N despite individual variability. **(F–H)** Startle amplitude across intensities (A.U.): **(F)** sexes pooled; **(G)** males, higher amplitude only for the repeated group at 82 dB; **(H)** females, higher amplitude in several LSD conditions across intensities. **(I)** Open-field examples: single-animal path traces. **(J)** Group-mean occupancy heatmaps (50×50 grid). **(K)** Total rotations by group (negative-binomial GLM, omnibus χ² = 8.02, P = 0.018); back-transformed means ± SE: Control 54.2 ± 12.6, Acute 136.2 ± 31.3, Repeated 81.4 ± 18.8; Tukey Control vs Acute P = 0.013. **(L)** Proportion of “high-rotators” (threshold = 95th percentile of Control; Fisher exact P = 0.43; n/N shown). **(M)** Occupancy centroids (x by y), sexes pooled; lines show group-wise linear fits with 95% confidence bands (R = 0.34, P = 0.079).

When sexes were combined, PPI was reduced after prenatal LSD (**Fig. 3C**). In sex-stratified analyses, males showed reduced PPI following repeated exposure (**Fig. 3D**), whereas females did not differ at the group level at this N despite clear individual variability (**Fig. 3E**). Startle amplitude was higher in females in both LSD conditions across intensities (**Fig. 3H**), and in males differed only for the repeated group at the highest intensity (**Fig. 3G**). Startle latency showed no consistent group effect. The only reliable change was a slower response at 82 dB in acute LSD females (**Fig. S2A-C**). These patterns indicate that amplitude changes do not account for the PPI reduction, which was most robust in males.

Open field revealed a stereotypy-dominant phenotype rather than gross hyperactivity. Representative single-mouse occupancy heatmaps are shown in **Fig. 3I**, with group-mean occupancy heatmaps in **Fig. 3J**. Total rotations were increased following prenatal LSD (negative-binomial GLM, Group LR χ²=8.02, P=0.018; back-transformed means ±SE: control 54.2 ±12.6, acute 136.2 ±31.3, repeated 81.4 ±18.8; Tukey control vs acute P=0.013; **Fig. 3K**). Using a control-anchored threshold (95th percentile of control), “high-rotators” were more frequent in LSD groups (acute 2/10, repeated 4/10, control 1/10; **Fig. 3L**). Spatial organization shifted without large changes in distance or velocity, as observed in pooled centroid distributions (**Fig. 3M**) and in the full set of raw path traces (**Fig. S2E**). Direction-specific rotation counts are provided in **Fig. S2F**.

Attempts to assess higher-order behaviors, including novel object recognition and social approach, were confounded by elevated baseline stereotypy and abnormal exploration, rendering these assays uninterpretable. Beginning at P21 and persisting through at least P156, 7 of 10 mice with repeated exposure and 5 of 10 with acute exposure exhibited continuous stereotypies (backflipping, tight circling, upright corner-hopping). At P156, females had lower body weight following repeated exposure (35.3 vs 43.7 g, P=0.021) and higher brain mass after acute exposure (0.635 vs 0.550 g, P=0.0001); males did not differ, and the acute female brain-mass effect remained significant after adjusting for body weight (**Fig. S3A-H**). Testis mass was unchanged; the testis-to-body ratio was higher after acute exposure (P=0.021). A blinded midline screen of atlas-matched coronal sections at P156 identified ventriculomegaly with frontal-horn predominance in LSD groups (control 1/10, acute 9/10, repeated 8/10; **Fig. S3I**) and rarer callosal variants, including asymmetric hypoplasia, an occasional supracallosal diverticulum continuous with the frontal horn, and partial agenesis of the anterior corpus callosum; given limited atlas levels and modest N, these reads are descriptive.

Overall, prenatal LSD exposure was associated with adult behavioral differences that were most robust in male offspring after repeated exposure (reduced PPI) and across groups as increased rotational stereotypy. Effect sizes for PPI were modest, and several higher-order assays were not interpretable due to elevated baseline stereotypy, which we therefore report as an important phenotype in its own right. Because we did not test persistence of the P8 cortical marker differences into adulthood or manipulate candidate receptors, we treat links between early cortical signatures and adult behavior as hypotheses motivated by the timing of embryonic exposure and the observed developmental phenotypes.

### Maternal serotonergic agonists activate embryonic choroid plexus and change CSF

Given the rapid appearance of LSD in embryonic CSF, we asked whether the embryonic ChP mounts an acute response to maternal serotonergic agonists. We quantified Fos induction after maternal dosing with LSD (0.3 mg kg⁻¹), psilocybin (3 mg kg⁻¹), 5-MeO-DMT (50 mg kg⁻¹), or the selective 5-HT₂C agonist WAY-161503 (3 mg kg⁻¹). In separate cohorts, we quantified epithelial apical remodeling and embryonic CSF protein after maternal dosing with LSD and the serotonergic psychedelics psilocybin and 5-MeO-DMT (**Fig. 4**).

**Fig. 4.**
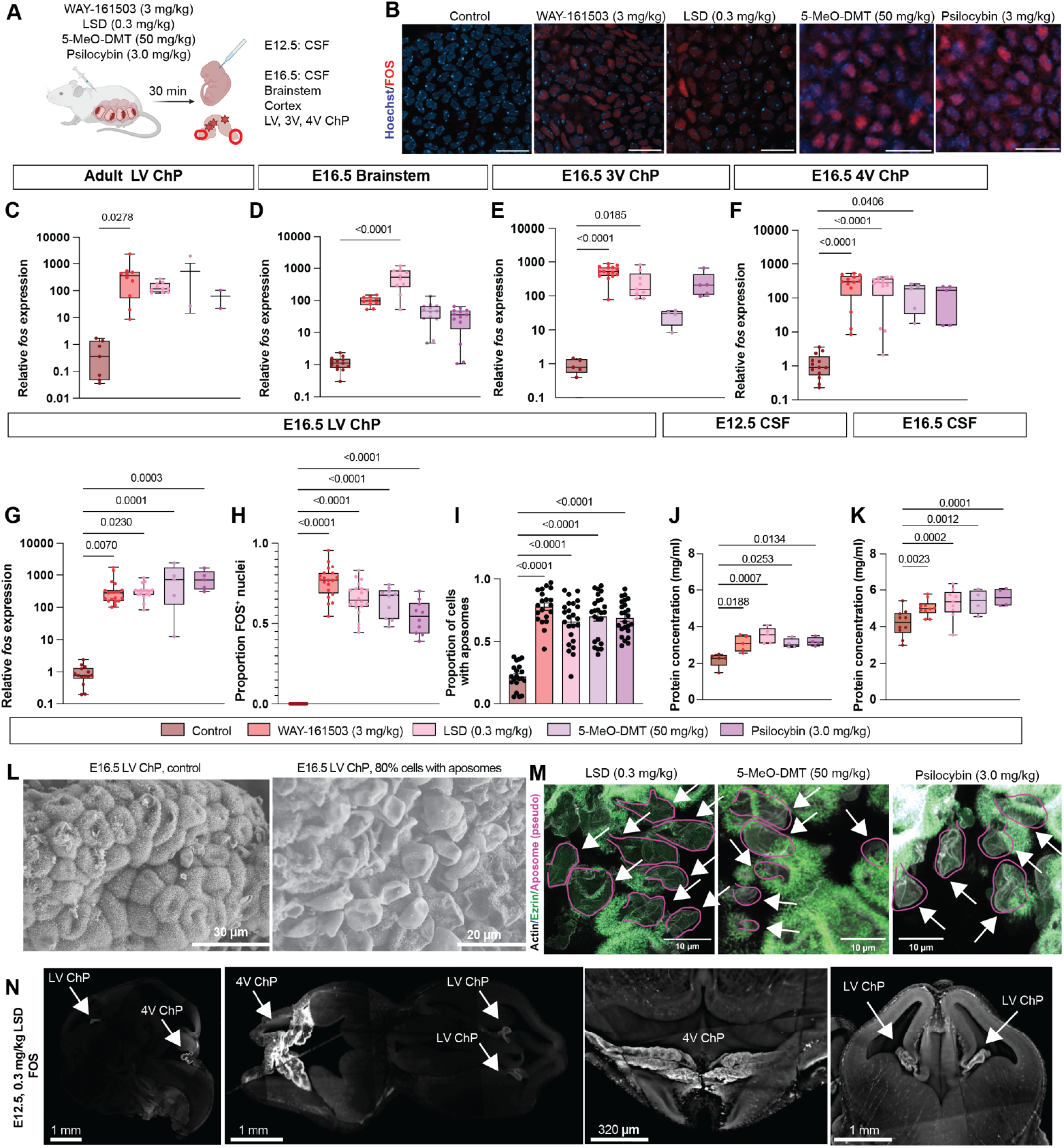
Maternal serotonergic agonists rapidly activate embryonic choroid plexus and increase embryonic CSF protein. **(A)** Experimental design: pregnant dams dosed at E12.5 (LSD 0.3 mg kg⁻¹; 5-MeO-DMT 50 mg kg⁻¹; psilocybin 3 mg kg⁻¹; WAY-161503 3 mg kg⁻¹); tissues and CSF collected 30 min later. Additional cohorts at E16.5 as indicated. **(B)** c-FOS immunostaining in ChP epithelium 30 min post-dose (representative fields; Hoechst/c-FOS). **(C)** Dam LV ChP: *Fos* expression (qPCR; 2^-ΔΔCt; normalized to *18s*, relative to vehicle). Control n = 7 ChP tissues, WAY n = 10, LSD n = 10, 5-MeO-DMT n = 2, psilocybin n = 2. **(D–F)** E16.5 ChP/brainstem: relative *Fos* expression in **D**, brainstem (Control n = 13 ChP tissues from 3 litters, WAY n = 10 from 3 litters, LSD n = 10 from 3 litters, 5-MeO-DMT n = 13 from 3 litters, psilocybin n = 13 from 3 litters); **E**, 3V ChP (Control n = 5 ChP tissues from 3 litters, WAY n = 15 from 3 litters, LSD n = 9 from 3 litters, 5-MeO-DMT n = 4 from 3 litters, psilocybin n = 5 from 3 litters); **F**, 4V ChP (Control n = 13 ChP tissues from 3 litters, WAY n = 13 from 3 litters, LSD n = 13 from 3 litters, 5-MeO-DMT n = 5 from 3 litters, psilocybin n = 5 from 3 litters). **(G)** E16.5 LV ChP: relative *Fos* expression (Control n = 14 ChP tissues from 3 litters, WAY n = 20 from 3 litters, LSD n = 17 from 3 litters, 5-MeO-DMT n = 5 from 3 litters, psilocybin n = 5 from 3 litters). **(H)** Proportion of FOS⁺ nuclei in E16.5 4V ChP (image-based quantification; control n = 13 ChP tissues from 3 litters, WAY n = 20 from 3 litters, LSD n = 20 from 3 litters, 5-MeO-DMT n = 10 from 3 litters, psilocybin n = 10 from 3 litters). **(I)** Proportion of ChP epithelial cells with apocrine blebs at 30 min (control n = 20 ChP tissues from 3 litters, WAY n = 20 from 3 litters, LSD n = 24 from 3 litters, 5-MeO-DMT n = 24 from 3 litters, psilocybin n = 24 from 3 litters). **(J–K)** Total protein concentration in embryonic CSF: **J**, E12.5 (control n = 5 pooled litters, WAY n = 5 pooled litters, LSD n = 5 pooled litters, 5-MeO-DMT n = 4 pooled litters, psilocybin n = 4 pooled litters); **K**, E16.5 (control n = 10 pooled litters, WAY n = 10 pooled litters, LSD n = 9 pooled litters, 5-MeO-DMT n = 4 pooled litters, psilocybin n = 4 pooled litters). (**L**) SEM of E16.5 LV choroid plexus comparing baseline aposome formation (left) with a high-frequency state (right; ∼80% aposome-positive epithelial cells). (**M**) High-resolution confocal images of E16.5 ChP epithelium 30 min after LSD, 5-MeO-DMT, or psilocybin exposure shows extensive apical blebbing and redistribution of actin and ezrin, a marker of microvilli and membrane curvature. **(N)** Light-sheet fluorescence imaging of optically cleared E12.5 heads stained for c-FOS 30 min post maternal 0.3 mg kg⁻¹ LSD (representative embryo). Cross-sections of whole-head dataset highlight strong activation in 4V ChP and detectable signal in LV ChP (arrows). Scattered bright foci are non-cellular artifacts (e.g., vascular-associated antibody retention and secondary antibody pooling) characteristic of this non-perfused, whole-embryo staining method. Scale bars as indicated.

A single maternal dose of LSD at E12.5 or E16.5 induced rapid activation of ChP epithelium across ventricles. The immediate early gene *Fos* was upregulated within 30 min, as measured by qPCR from microdissected ChP collected 30 min after dosing in adult dams (LV ChP) and in E16.5 embryos (LV/3V/4V ChP and brainstem; **Fig. 4C–G**) and by c-FOS immunostaining (**Fig. 4B,H**). The response generalized to WAY-161503, 5-MeO-DMT, and psilocybin (**Fig. 4D–F,G**). Whole-head light-sheet imaging of optically cleared E12.5 embryos further confirmed robust c-FOS in 4V ChP with detectable, but less intense signal in LV ChP (**Fig. 4L, Fig. S4A-F**). Additional FOS signal was observed in the locus coeruleus in these datasets (anatomical observation; not quantified; **Fig. S4G–H**).

Downstream of transcriptional activation, ChP cells showed rapid apical remodeling. Within 30 min of maternal LSD, psilocybin, or 5-MeO-DMT, epithelial cells exhibited increased aposome formation (actin-rich, secretory protrusions [phalloidin/ezrin]), consistent with an apocrine-like secretory state (Courtney and Lehtinen, 2025; Courtney et al., 2025) (**Fig. 4B,I,L,M**). Functionally, total protein concentration rose in embryonic CSF 30 min following maternal psychedelics at both E12.5 and E16.5 (**Fig. 4J–K**). Together, these data support that maternal psychedelic exposure activates embryonic ChP leading to rapid transcriptional, cytoskeletal, and secretory responses that alter CSF composition.

## Discussion

Maternally administered LSD rapidly reached embryonic CSF during mid-gestation and coincided with an acute response in embryonic ChP epithelium, including Fos induction, apical remodeling, and increased CSF protein. In separate cohorts, prenatal exposure during E12.5–E16.5 was associated with altered S1 cellularity and projection-neuron subtype marker composition at P8, and with adult behavioral differences most evident as increased rotational stereotypy and modest, male-predominant reductions in prepulse inhibition after repeated exposure. Together, these findings position the embryonic CSF–choroid plexus interface as a rapidly engaged compartment through which maternal serotonergic agonists can access and influence embryonic signaling environments.

A key implication of the pharmacokinetic data is that embryonic CSF exposure occurs on a timescale compatible with acute epithelial and neural responses. The ChP is well-positioned to respond to CSF-borne ligands, and transcriptomic atlases and prior *in vitro* work suggest serotonergic responsiveness in ChP epithelium, including prominent *Htr2c* expression. Consistent with this, we observed rapid Fos induction across ventricles after maternal dosing with LSD and with other serotonergic agonists, including a selective 5-HT_2_C agonist. We interpret these data as supporting a model in which serotonergic agonists can acutely engage ChP epithelium during development. However, our study did not test receptor necessity. LSD binds serotonergic sites across multiple brain regions, though ChP epithelium harbors the highest reported site density (Yagaloff and Hartig, 1985); resolving the relative contributions of ChP-mediated versus direct neural actions, including at 5-HT_2_A-expressing neuronal populations, and potential effects of active metabolites, will require targeted perturbation experiments.

The early postnatal cortical signatures observed at P8 likely reflect a combination of processes – altered progenitor behavior, maturation state, survival, or subtype specification – that shift the relative abundance of projection-neuron subtype marker populations. We used SATB2, CTIP2, and TBR1 as operational indices of major projection-neuron subclasses during early postnatal development; future work will be needed to determine whether these differences reflect fate specification versus later developmental remodeling and whether they persist into adulthood. Increased microglial density and astrocytic reactivity in exposed offspring suggest a broader developmental response that may interact with neuronal maturation, but mechanistic relationships among these findings remain to be determined.

Behaviorally, prenatal LSD exposure was associated with increased stereotyped rotation and with PPI reductions that were most robust in males after repeated exposure. These outcomes are not specific to any one neurodevelopmental condition, and we therefore interpret them as behavioral readouts consistent with altered circuit function rather than as disease models. Taken together, our data support a temporally coherent model in which rapid embryonic CSF access and ChP activation coincide with downstream cortical and behavioral phenotypes. Because the embryonic ChP assays, P8 histology, and adult behavior were collected in separate cohorts and without receptor/ChP perturbation, we present the ChP→cortex→behavior sequence as a working model rather than a demonstrated causal pathway.

Several limitations are important for interpretation. First, we used a single dose level and focused on acute versus repeated exposure regimens, which constrains inferences about dose-response relationships. Second, our dose selection was guided by rodent literature (Halberstadt and Geyer, 2013; Maitland et al., 2025; McGriff et al., 2025) and the need for robust exposure and assay sensitivity in embryonic CSF; it is not intended to model human exposure levels, and we do not attempt to estimate pregnancy risk. Addressing these limitations with PK/PD-anchored dosing, receptor perturbation (e.g. pharmacological antagonism or genetic approaches), and longitudinal anatomical/behavioral mapping will be essential for resolving causal mechanisms.

In summary, our data show that maternal LSD can access embryonic CSF on a minutes timescale and that embryonic ChP epithelium mounts a rapid response to serotonergic agonists during cortical development. These findings provide a framework for studying how maternal serotonergic exposures may engage the embryonic CSF-choroid plexus interface and motivate future mechanistic experiments to define receptor pathways and downstream developmental consequences.

## Acknowledgments

We thank the NIDA Drug Supply Program for providing controlled substances (LSD, psilocybin, and 5-MeO-DMT) used in this study. We are grateful to Chinfei Chen, Hisashi Umemori, and Harry Cramer for imaging advice and support, and to the BCH IDDRC Cellular Imaging Core (<u>RRID: SCR_026485)</u> for confocal and light sheet imaging. We are especially grateful to the Boston Children’s Hospital Research Controlled Substances Team for their guidance throughout this project. We thank Nathaniel Hodgson and the BCH Animal Behavior and Physiology Core for assistance with behavioral assays. We thank Sofia Bennetts and Pura Arroyo for technical support and help with figures. We are grateful to syGlass (SyGlass, Inc.) for extended access to their visualization software under a trial license and for technical assistance. syGlass had no role in study design, data collection, data analysis, or the decision to publish.

Figures were created in part using BioRender.com. We thank members of the Lehtinen lab for valuable feedback and discussions. This work was supported by the National Institutes of Health (R01NS088566 to M.K.L.; P50HD105351 and S10OD030322 to the BCH IDDRC), the Howard Hughes Medical Institute Gilliam Fellowship (Y.C.), the Jane Coffin Childs Memorial Fund for Medical Research Fellowship (Y.C.), and the Kirby Innovation Award (M.K.L.).

## Author contributions

Conceptualization: YC.

Methodology: YC, JMA.

Investigation: YC, JMA, CLL.

Visualization: YC, JMA.

Funding acquisition: YC, CJW, MKL.

Project administration: CJW, MKL.

Supervision: CJW, MKL.

Writing – original draft: YC.

Writing – review & editing: YC, JMA, CLL, CJW, MKL.

## Data and materials availability

All data needed to evaluate the conclusions of this study are present in the paper and/or the Supplementary Materials. Additional data and analysis scripts are available from the corresponding author upon reasonable request. Controlled substances (LSD, psilocybin, 5-MeO-DMT) were obtained from the NIDA Drug Supply Program and cannot be redistributed; qualified investigators should request these materials directly from NIDA.

## References

1. Bonnin A, Goeden N, Chen K, Wilson ML, King J, Shih JC, Blakely RD, Deneris ES, Levitt P (2011) A transient placental source of serotonin for the fetal forebrain. Nature:2–7.

2. Carvajal-Oliveros A, Campusano JM (2021) Studying the Contribution of Serotonin to Neurodevelopmental Disorders. Can This Fly? Front Behav Neurosci 14:601449.

3. Courtney Y, Head JP, Dani N, Chechneva OV, Shipley FB, Zhang Y, Holtzman MJ, Sadegh C, Libermann TA, Lehtinen MK (2025) Choroid plexus apocrine secretion shapes CSF proteome during mouse brain development. Nat Neurosci:1–14.

4. Courtney Y, Lehtinen MK (2025) Apocrine secretion by the choroid plexus. Fluids and Barriers of the CNS 22:75.

5. Damkier HH, Brown PD, Praetorius J (2013) Cerebrospinal fluid secretion by the choroid plexus. Physiol Rev 93:1847–1892.

6. Dani N et al. (2021) A cellular and spatial map of the choroid plexus across brain ventricles and ages. Cell 184:3056–3074.e21.

7. Fame RM, Lehtinen MK (2020) Emergence and Developmental Roles of the Cerebrospinal Fluid System. Dev Cell 52:261–275.

8. Gaspar P, Cases O, Maroteaux L (2003) THE DEVELOPMENTAL ROLE OF SEROTONIN : NEWS FROM MOUSE MOLECULAR GENETICS. Nature Reviews Neuroscience 4.

9. Gato A, Desmond ME (2009) Why the embryo still matters: CSF and the neuroepithelium as interdependent regulators of embryonic brain growth, morphogenesis and histiogenesis. Developmental Biology 327:263–272.

10. Halberstadt AL, Geyer MA (2013) Characterization of the head-twitch response induced by hallucinogens in mice: detection of the behavior based on the dynamics of head movement. Psychopharmacology (Berl) 227:10.1007/s00213-013-3006-z.

11. Lehtinen MK, Zappaterra MW, Chen X, Yang YJ, Hill AD, Lun M, Maynard T, Gonzalez D, Kim S, Ye P, D’Ercole AJ, Wong ET, LaMantia AS, Walsh CA (2011) The Cerebrospinal Fluid Provides a Proliferative Niche for Neural Progenitor Cells. Neuron 69:893–905.

12. Lun MP, Monuki ES, Lehtinen MK (2015) Development and functions of the choroid plexus-cerebrospinal fluid system. Nature Reviews Neuroscience 16:445–457.

13. MacAulay N, Keep RF, Zeuthen T (2022) Cerebrospinal fluid production by the choroid plexus: a century of barrier research revisited. Fluids Barriers CNS 19:26.

14. Maitland AD, Gonzalez NR, Walther D, Pereira F, Baumann MH, Glatfelter GC (2025) Rapid, Open-Source, and Automated Quantification of the Head Twitch Response in C57BL/6J

15. Mice Using DeepLabCut and Simple Behavioral Analysis. ACS Pharmacol Transl Sci 8:2694–2709.

16. McGriff SA, Hecker JC, Maitland AD, Partilla JS, Baumann MH, Glatfelter GC (2025) Psychedelic-like effects induced by 2,5-dimethoxy-4-iodoamphetamine, lysergic acid diethylamide, and psilocybin in male and female C57BL/6J mice. Psychopharmacology Available at: 10.1007/s00213-025-06795-x [Accessed August 17, 2025].

17. Sanders-Bush E, Breeding M (1991) Choroid plexus epithelial cells in primary culture: a model of 5HT1C receptor activation by hallucinogenic drugs. Psychopharmacology (Berl) 105:340–346.

18. Saunders NR, Dziegielewska KM, Fame RM, Lehtinen MK, Liddelow SA (2023) The choroid plexus: a missing link in our understanding of brain development and function. Physiological Reviews 103:919–956.

19. Shah R, Courtiol E, Castellanos FX (2018) Abnormal Serotonin Levels During Perinatal Development Lead to Behavioral Deficits in Adulthood. Frontiers in Behavioral Neuroscience 12:1–10.

20. Yagaloff KA, Hartig PR (1985) 125I-lysergic acid diethylamide binds to a novel serotonergic site on rat choroid plexus epithelial cells. J Neurosci 5:3178–3183.

21. Zappaterra MW, LaMantia AS, Walsh CA, Lehtinen MK (2013) Isolation of Cerebrospinal Fluid from Rodent Embryos for use with Dissected Cerebral Cortical Explants. J Vis Exp:50333.

