## Supplementary Information for "Gestational LSD exposure in mouse rapidly reaches embryonic CSF and is associated with altered choroid plexus signaling, cerebral cortical development, and offspring behavior"

### Materials and Methods for

**Affiliations:**

#### Materials and Methods

Ethical regulations. Our research complies with all relevant ethical regulations. The Boston Children's Hospital IACUC approved all mouse studies as per protocols 00002094 and 00002095.

Materials.

Antibodies.

- AQP1 Rabbit polyclonal; IHC 1:250 Millipore Cat#AB2219; RRID AB_1163380
- C-FOS Rabbit monoclonal recombinant IgG Clone Rb108B5; IHC 1:1000 Synaptic Systems Cat#226 008; RRID AB_2891278
- C-FOS Mouse monoclonal IgG1 Clone E-8; IHC 1:50 Santa Cruz Cat#E-8 SC-166940; RRID AB_10609634
- CTIP2 Rat monoclonal Clone 25B6; IHC 1:250 Abcam Cat#ab18465; RRID AB_2064130
- Ezrin Mouse monoclonal; IHC 1:100 Invitrogen Cat#35-7300
- F-actin Phalloidin-iFluor 647; IHC 1:1000 Abcam Cat#ab175759
- GFAP Chicken polyclonal; IHC 1:200 Abcam Cat#ab4674; RRID AB_304558
- IBA1 Goat polyclonal; IHC 1:200 Abcam Cat#ab5076; RRID AB_91676
- PV (Parvalbumin) Guinea pig polyclonal IHC 1:250 Synaptic Systems Cat#195 004
- SATB2 Mouse Monoclonal; Clone 2E6 IHC 1:100 Abcam Cat#ab51502; RRID AB_882455
- TBR1 Rabbit polyclonal; IHC 1:1000 Abcam Cat#ab31940; RRID AB_2200219
- Donkey anti Goat Alexa fluor 488; IHC 1:250 Abcam Cat#AB96931; RRID AB_10680169

- Goat anti Mouse Alexa fluor 594; IHC 1:250 Life Technologies Cat#A-11005; RRID AB_2534073
- Goat anti Mouse Alexa fluor 647; IHC 1:250 Thermo Fisher Cat#A-21235; RRID AB_2535804
- Goat anti Rabbit Alexa Fluor 488; IHC 1:250 Life Technologies Cat#A11034; RRID AB_2758380
- Goat anti Rabbit Alexa Fluor 594; IHC 1:250 Abcam Cat#a11012; RRID AB_2534079
- Goat anti Rat Alexa fluor 568; IHC 1:250 Life Technologies Cat#A-11077; RRID AB_2534121

<u>Chemicals, Peptides, and Recombinant Proteins.</u>

- WAY-161503 (Tocris, Cat#1801)
- (+)-lysergic acid diethylamide (+)-tartrate (NIDA Drug Supply Program, Material #7315-004)
- Psilocybin (NIDA Drug Supply Program, Material #7437-001)
- 5-methoxy-N,N-dimethyltryptamine (5-MeO-DMT) (NIDA Drug Supply Program, Material #7431-001)
- Sterile 0.9% sodium chloride (USP)
- d3-LSD analytical standard, >99% (Cerilliant, Round Rock, TX) — used as internal standard
- LSD analytical standard, >99% (Cerilliant)
- Acetonitrile, methanol, water, formic acid, LC-MS grade (Fisher Scientific, Optima)

- Artificial CSF (Tocris)
- Blank CD1 mouse serum (Innovative Research, Novi, MI)

Consumables

- Fluoromount-G, Thermo Fisher Scientific, Cat# 00-4958-02
- Hoechst 33342 (Molecular Probes™ Hoechst 33342, Trihydrochloride, Trihydrate Cat#H3570)
- LifeCanvas Passive Clearing Kit Cat#C-PCK-250: SHIELD Epoxy, SHIELD Buffer, SHIELD ON, Delipidation Buffer, EasyIndex
- OCT (Optimal Cutting Temperature) Compound
- Phenomenex Kinetex Phenyl-Hexyl column, 1.7 µm, 100 × 1.2 mm, 35 °C used
- Sodium Azide, Sigma-Aldrich, Cat# S2002
- Sucrose, Sigma-Aldrich, Cat# S0389
- Triton X-100, Sigma-Aldrich, Cat# T8787
- Waters Sirocco Protein Precipitation Plates

Equipment

- Sciex QTRAP 5500 mass spectrometer
- Waters ACQUITY UPLC I-Class system
- Waters Positive Pressure-96 Manifold
- Evaporex EVK-96 evaporator
- MetaQuest VR headset (for light sheet figure renderings)
- Roche LightCycler 480II
- Thermo Scientific NanoDrop 1000

- Zeiss LSM 710, 20× air; 63× oil objectives
- Zeiss LSM 980, 20× air; 63× oil objectives
- Zeiss Lightsheet 7, 5x objective

Mice. Animals were housed in a temperature- and humidity-controlled room (21 ± 1.5°C, 35–70% humidity) on a 12 h light/12 h dark cycle (7 a.m. on 7 p.m. off) and had free access to food and water. We used equal numbers of male and female mice unless otherwise stated. Adult and timed-pregnant CD-1 (RRID:MGI:5649524) mice were obtained from Charles River Laboratories, Jackson Laboratory, or bred in-house.

Oligonucleotides

- Mm00487425_m1; Fos, FAM-MGB. ThermoFisher. cat#4331182
- Hs99999901_s1; Eukaryotic 18s rRNA, VIC- MGB. ThermoFisher, cat#4331182

Software.

- Olympus FV31S-SQ (FluoView, version 2.6)
- Noldus EthoVision XT videotracking software (version 9.0, Noldus Information Technologies, Leesburg, VA)
- Avisoft Recorder (version 2.97; Avisoft Bioacoustics)
- Sonotrack Call Classification Software (version 1.4.7, Metris B.V., Netherlands).
- Observer XT software (version 9.0, Noldus)
- R versions 3.6.3 and 4.1.2
- Biodock.AI
- ImageJ/FIJI Version 2.9.0/1.55i

- Ingenuity Pathway Analysis (QIAGEN Inc.)
- GraphPad Prism 9 and 10
- QuPath Version 0.5.0
- Ggplot2 Version 3.4.0
- Dplyr Version 1.1.0
- Tidyr Version 1.3.0
- Zen Black 3.6
- Zen Blue 3.7
- Sciex Analyst (for acquisition)
- Waters ACQUITY Console (for LC control)
- Sciex MultiQuant v3.0.3 (for quantitation)
- syGlass v2.3.0

#### Methods

##### Behavioral analyses.

Behavioral analyses were carried out on CD-1 mice, 10 mice per condition beginning at age P90, in collaboration with the Boston Children's Hospital Animal Behavior & Physiology Core.

Open-Field Test. Adult CD-1 mice (n=10 per condition unless noted) explored a 40×40 cm arena for 15 min. Primary outcomes were total distance traveled and rotational stereotypy; spatial occupancy metrics (e.g., centroid/heatmaps) were used as descriptive visualizations. Rotations were quantified from the animal's instantaneous heading angle relative to the arena center. A “rotation” was defined as ≥360° cumulative angular displacement in a single direction, with direction determined by the sign of angular velocity. General Considerations. Several precautions were taken to ensure the accuracy and reliability of the test results. Investigators who owned cats or ferrets were required to wear clothing that had not come into contact with these pets on the day of testing, as the scent of predators could elicit strong fear responses in mice. Mice were placed in the testing room at least 30 minutes before the start of the experiment to acclimate to the environment. The experimenter remained blind to the conditions of the test mice, and mice were randomly assigned to experimental groups, with control groups consisting of littermates whenever possible. The test arena was thoroughly cleaned between trials by removing feces and urine, spraying with Clidox, and wiping with a water-dampened paper towel to eliminate any residual odors from previous mice. Each mouse was only tested once to avoid the confounding effects of prior exposure on subsequent behavior.

Procedure. The Motor Monitor software was used to track and record the movements of the mice during the test. The current configuration was set to "OFHB Single 9 Hole," which activated the bottom beam for locomotion tracking. For each testing session, parameters such as test duration, pre-pause duration, and mouse IDs were manually entered or pre-set using a template to streamline the process.

Testing began by placing each mouse in the arena, with sessions initiated either by the mouse's movement (Start Session on X-Y Activity) or manually by the experimenter (Start Session by Start Button Only). The session was automatically timed, and data were saved to a session file for later analysis. After each session, the mouse was returned to its home cage, and the arena was cleaned in preparation for the next test.

Data Reduction. Data from the OFT sessions were processed using the Motor Monitor software. The session files were selected and reduced into a single Excel file, with specific parameters chosen for analysis, such as distance traveled, time in the center and periphery, and rearing bouts. The processed data were saved as a .csv file for further statistical analysis.

Pre-Pulse Inhibition.

Pre-pulse inhibition (PPI) was assessed to measure sensorimotor gating in the test subjects. PPI occurs when a weak pre-stimulus (pre-pulse) reduces the response to a subsequent stronger startle stimulus. This phenomenon is an important indicator of an organism's ability to filter out unnecessary information and has been associated with various neuropsychiatric conditions such as schizophrenia and Alzheimer's disease. While PPI is not diagnostic of a particular disorder, it provides valuable insight into the functionality of underlying neural circuits and neurotransmitter systems.

General Considerations. To ensure accurate results, several precautions were taken. Investigators who owned cats or ferrets wore clothing that had not come into contact with these pets on the day of the experiment, as the scent of these animals can trigger a strong fear response in mice due to predator cues. Mice were placed in the testing room at least 30 minutes before the start of the experiment to acclimate to the environment. Throughout the experiment, the experimenter remained blind to the conditions of the test mice, and test mice were randomized, with control groups consisting of littermates wherever possible. Between trials, the test chambers were thoroughly cleaned by removing any feces and urine, spraying with Clidox, and wiping with a water-dampened paper towel to eliminate any residual odors from previous mice.

Procedure. PPI was evaluated using a protocol that varied the intensity of the pre-pulse while keeping the interval between the pre-pulse and the startle pulse constant. The testing system was pre-programmed with the specific experimental settings. The auditory pre-pulse levels varied between 70, 74, 78, and 82 dB, each lasting 20 milliseconds. The background noise in the testing chambers was maintained at a constant 60 dB throughout the experiment. The interval between trials varied randomly between 10 and 30 seconds to reduce the predictability of the stimulus. The startle pulse, set at 105 dB and lasting 40 milliseconds, was delivered 100 milliseconds after the pre-pulse. Testing was conducted with four mice at a time, with a recommended sample size of eight mice per group. Each mouse was given a five-minute acclimation period in the test chamber, where they were exposed to 60 dB background noise. During the testing phase, each mouse underwent 20 trials in total, with four pre-pulse levels presented in random order, repeated five times.

Data Analysis. PPI was calculated using the formula: %PPI = 100 × [(pulse-alone) – (pre-pulse + pulse score)] / pulse-alone score. This formula quantifies the percentage of inhibition of the startle response when a pre-pulse is presented, providing a measure of sensorimotor gating in the mice.

ChP dissection.

Whole embryonic and adult ChP tissues were isolated as follows. LV ChP: Cerebral cortical hemispheres were separated. Following incisions into the cortical tissue at either end of the hippocampus, the hippocampus was rolled out using the flat surface of a microscalpel and the underlying LV ChP was separated from the hippocampus/fornix. 3V ChP: The dorsal midline of the midbrain was exposed and 3V ChP was isolated. Specific to adult brain, flushing the midline region with 1xHBSS helped visualize the ChP, which was identifiable by a blood vessel running along its rostro-caudal axis. The adult 3V ChP extends ventrally and rostrally, ultimately connecting to the ventral base of the LV ChP. 4V ChP: Hindbrain was separated from the brain.

The cisterna magna was exposed by gently guiding away the developing or mature cerebellum to expose the 4V ChP.

In vivo drug delivery.

All agonists were injected subcutaneously into timed-pregnant CD1 female mice in 0.9% sterile saline. Doses were: WAY-161503 3 mg $kg^{-1}$ (Tocris, Cat#1801); (+)-lysergic acid diethylamide tartrate 0.3 mg $kg^{-1}$ (NIDA Drug Supply Program); psilocybin 3 mg $kg^{-1}$ (NIDA Drug Supply Program); 5-methoxy-N,N-dimethyltryptamine (5-MeO-DMT) 50 mg $kg^{-1}$ (NIDA Drug Supply Program). WAY-161503 was delivered as a 10 mM solution at 1 µL × body weight (g). (+)-lysergic acid diethylamide tartrate was delivered as a 0.6 mM solution at 0.6 µL × g. Psilocybin was delivered from a 15.03 mg $mL^{-1}$ working solution at 0.1995 µL × g; when the calculated drug volume was <50 µL, sterile saline was added to a total injection volume of 50 µL without altering the psilocybin amount. 5-MeO-DMT was delivered from a 50 mg $mL^{-1}$ working solution at 1 µL × g. For each dam, body weight was recorded immediately prior to dosing, and injection time, time of sacrifice, dissection completion, and litter size were logged.

Image acquisition and quantification.

Images were acquired with a Zeiss LSM710 or LSM980 microscope at the Boston Children's Hospital Cellular Imaging Core. Images of ChP explants and brain sections were acquired using a 20x air or 63x oil objective. Images of ChP explants prepared with protein-retention expansion microscopy were acquired with a 63x NA1.2 water-immersion objective. ZEN Black software was used for image acquisition and ZEN Blue used for Airy processing.

Aposome quantification. Aposome quantification was performed by author YC while blinded to treatment condition (images coded prior to scoring). YC took 4 20x fields of view per ChP explant and counted epithelial cells displaying apocrine structures via either SEM or immunostaining. The proportion of cells with apocrine structures was quantified as the number of epithelial cells with an apocrine protrusion divided by the total number of visible epithelial cells. The proportion of cells with apocrine structures for each of the 4 fields of view was then averaged and used as N=1 biological replicate per embryo.

C-FOS/FOS positive nucleus quantification. FOS quantification was performed with ImageJ/FIJI. 4 20x fields of view were taken per ChP explant, and the average proportion of FOS positive nuclei in these 4 fields of view was used as N=1 biological replicate per CD-1 embryo. Each field of view was thresholded, then the "Analyze particles" function was used with a size discrimination of over 200 px. Counting was performed for FOS channels and DAPI channels, and then the number of FOS nuclei was divided by the total number of nuclei to obtain the "Proportion of FOS positive nuclei."

Excitatory neuronal marker counting and co-expression analysis. Cerebral cortical layer marker counting and colocalization was performed using Biodock software [1] for CD-1 mice. A fully automated AI model was trained by a 3-step iterative process on 3 test staining images containing CTIP2, SATB2, TBR1, and DAPI. Input images were 4 channels. TIF files obtained by importing the .CZI file from the microscope into FIJI, rotating, and cropping a rectangular region comprising the primary somatosensory cortex from just above layer 1 to the corpus callosum.

The model was trained to classify each cell in the cerebral cortex into one of 8 classes based on co-expression of each marker used. Potential cell classes were:

1) CTIP2- SATB2- TBR1- (DAPI only)

2) CTIP2+, SATB2-, TBR1-

3) CTIP2+, SATB2+, TBR1-

4) CTIP2+, SATB2+, TBR1+

5) CTIP2+, SATB2-, TBR1+

6) CTIP2-, SATB2+, TBR1-

7) CTIP2-, SATB2+, TBR1+

8) CTIP2-, SATB2-, TBR1+

Each image's AI analysis was carefully checked by author YC, validating the correct identification of cells and visualizing overall distribution of labels.

The data resulting from the Biodock AI model is a .CSV file containing a comprehensive list of cell measurements obtained from cortical tissue samples. The data included several parameters for each cell: Object ID, Image Origin, Class, Area, Eccentricity, Length of Major and Minor Axes, Perimeter, Solidity, X and Y Positions, and Average Intensity across four channels.

Further analysis was conducted in the R programming environment. We utilized key packages including ggplot2 [2] for data visualization, dplyr [3] for data manipulation, and tidyr [4] for data tidying. A critical step in our analysis involved segmenting the cortical area into 6 equal-sized bins from top to bottom. To achieve this, we calculated the minimum and maximum values of the Y Position from our data, establishing six equally spaced bins along the Y-axis. This segmentation allowed us to analyze cell distribution across different cortical layers. Each cell was assigned to one of the six bins based on its Y Position, facilitating a layer-specific analysis of cell distribution. We then performed a count of cells in each bin, categorizing them into classes based on their expression of markers: CTIP2, SATB2, and TBR1.

We computed the percentage of each cell class in each bin, relative to the total cell count in that bin. Analyzed data were exported into a new .CSV file containing the expanded summary table with cell counts, percentages, and other relevant statistics. These data were imported into GraphPad Prism for statistical analyses and graphing. The R script used for this analysis is available at [5].

IBA1+ cell quantification. Images were acquired on a LSM980 microscope at the Boston Children's Hospital Cellular Imaging Core using a 20x air objective. Quantification was completed in FIJI. 100 µm by 100 µm regions of interest across the primary somatosensory cortex, corpus callosum, or hippocampus were acquired. IBA1+ cells were counted manually by YC as cells with staining for IBA1+ and DAPI. 3 fields of view were acquired from each of 3 technical replicate images from each P8 CD-1 mouse. Each value reported is an average of these 9 images per mouse.

Immunostaining. *Choroid plexus explants.* ChP explants were dissected from mice and fixed in 4% paraformaldehyde for 10 min at room temperature [6]. Samples were incubated in primary antibodies overnight at 4ºC, shaking and in secondary antibodies at room temperature for two hours, shaking. All antibodies were diluted in 0.1% Triton X-100 in PBS. No blocking or antigen retrieval methods were used. All samples were counterstained with Hoechst 33342 (Invitrogen, H3570, 1:10,000) and mounted onto slides using Fluoromount-G (SouthernBiotech).

*Brain sections.* Postnatal animals were perfused with PBS followed by 4% PFA. The brains were quickly dissected and post-fixed with 4% PFA overnight. Samples were cryoprotected and prepared for embedding at 4°C with 10% sucrose, 20% sucrose, 30% sucrose, 1:1 mixture of 30% sucrose and OCT (overnight), and OCT (1 h on ice). Samples were frozen in OCT. Antigen retrieval and blocking were not performed. Cryosections were permeabilized (0.1% Triton X-100 in PBS), incubated in primary antibodies at 4°C overnight and secondary antibodies for 2 h at room temperature. Hoechst 33,342 (Invitrogen H3570, 1:10,000, 5 min at room temperature) was used to visualize nuclei. Sections were mounted on glass slides with Fluoromount-G (SouthernBiotech).

Images were acquired using Zeiss LSM710 or LSM980 confocal microscope with 20× objective. ZEN Black software was used for image acquisition and ZEN Blue used for Airy processing.

Embryonic CSF collection (E12.5 and E16.5). Timed-pregnant CD-1 dams were euthanized and embryos were rapidly harvested under a stereomicroscope. Embryonic CSF was collected from the cisterna magna using a pulled glass microcapillary with a fine tip by gentle capillary action/aspiration under stereomicroscopic guidance. Samples were maintained on ice, then spun 1000× *g* for 10 min at 4 °C. The supernatant was collected and used for analysis. Samples with visible blood or tissue contamination were discarded. When required to obtain sufficient volume for downstream assays, CSF from multiple embryos within a litter was pooled to generate a single biological replicate (see figure legends for pooling scheme and n). Samples were stored at −80 °C and underwent a single freeze–thaw prior to assay.

Embryonic CSF total protein quantification (BCA). Total protein concentration in embryonic CSF was measured using the Pierce BCA Protein Assay Kit **(**Thermo Fisher Scientific) according to the manufacturer's microplate protocol. Standards and samples were run in duplicate**.** For each well, 1 µL CSF was assayed directly and mixed with 200 µL working reagent. Plates were incubated at 37 °C for 30 min, cooled to room temperature, and absorbance was read at 562 nm**.** Protein concentrations were interpolated from the standard curve and reported as mg/mL.

Pharmacokinetic Analysis via LC-MS/MS.

Pharmacokinetic analysis of serum and cerebrospinal fluid samples was performed on a Sciex 5500 Q-Trap MS coupled to a Waters Acquity UPLC system. Water and acetonitrile with 0.1% Formic Acid served as the solvent system, run at a flow rate of 0.375mls/min through a Kinetex 1.7um 100x1.2mm Phenyl-Hexyl column held at 35C. The LC Method consisted of a 5-

40% B Gradient for 1.75 min, followed by a 95%B wash step held until 2.5 minutes, system was returned to 5%B from 2.5 to 2.75 minutes and held at 5%B until 3 minutes. Samples were run in MRM, positive ionization mode. Unique transitions were monitored for both LSD and d3-LSD which served as the internal standard (ISTD; **Supplementary Table 1**). Source conditions are available in **Supplementary Table 2**.

Data was analyzed using MultiQuant software. Area under the curve (AUC) of the calibrators was adjusted in conjunction with the AUC of the ISTD, and used to establish a quadratic curve fit with 1/x2 weighting. All standard curves generated maintained $R^2$ values between 0.996-0.999, using only calibrators >85% accuracy.

Materials. Analytes d3-LSD (>99%) and LSD (>99%) were purchased from Cerilliant (Round Rock, Texas). Optima LC-MS grade acetonitrile, methanol, water and formic acid purchased from Fischer Scientific (Hampton, NH). Artificial CSF was purchased from Tocris Bioscience (Minneapolis, MN), and blank CD1 serum Innovative Research (Novi, MI). Sirocco Protein Precipitation Plates were acquired from Waters (Milford, MA).

Equipment. Samples were analyzed using a Sciex Q-TRAP 5500 Mass Spectrometer coupled to a Waters Acquity UPLC I-Class system. The mass spectrometer was managed using Sciex Analyst Software and the UPLC using the Waters Acquity Console. Sample processing was completed using a Waters Positive Pressure-96 Manifold and Evaporex EVK-96. Data was analyzed using Sciex MultiQuant (version 3.0.3).

Sample Processing. Standard curves were generated using Artificial CSF (aCSF) or blank serum as matrix, d3-LSD serving as the internal standard. Calibrators were prepared from methanol stocks of analytical standards, establishing a standard curve from 0.25ng/mL to 200ng/mL in aCSF and from 0.25ng/mL to 600ng/mL in blank serum.

Calibrators and samples were precipitated with Acetonitrile, then filtered using Waters Sirocco Protein Precipitation plates and a Waters Positive Pressure Manifold. The flow-through was dried down and resuspended in 5% Mobile Phase B and spun down to remove any precipitate.

Samples were injected in triplicate with blanks in between sample injections, sets submitted in a randomized fashion.

Quantification and statistical analysis.

To achieve robust and unbiased results, we performed our analyses across multiple samples, prospectively randomized treatment group via random number generation, quantified results in a double-blinded manner and performed rigorous pre- and post-hoc statistical analyses to test our hypotheses. We consulted with Harvard statisticians (BCH, BIDMC, Broad) to ensure proper calculations of power analyses and statistical methods, including accounting for multiple comparisons and covariates. We performed an equal number of experiments on male and female mice (except for pregnant mice which were all female) and carefully examined sex as a biological variable in our analyses. For each experiment, pilot studies were used to determine variance, and two-tailed power analyses determined optimal group size for alpha = 0.05 and beta = 0.80. Biological replicates (*N*) were defined as samples from individual animals, analyzed either in the same experiment or within multiple experiments, except when individual animals could not provide sufficient sample material (i.e., CSF), which required pooling samples from multiple animals into one biological replicate as described.

Statistical analyses were performed using Prism (GraphPad) v9 or R (version 4.1.2). Outliers were excluded using the ROUT method (Q = 1%). Data distribution was assessed using the Shapiro-Wilk normality test, and homogeneity of variances between datasets was evaluated using F tests or Bartlett's tests. Parametric tests were used when data were normally distributed, and variances were approximately equal; otherwise, nonparametric alternatives were applied. Analyses were blinded; ‘litter’ was tracked and used as a random intercept in sensitivity models.

Specific statistical tests were selected based on the data and analysis goals: Student’s two-tailed unpaired t test was used for comparisons with one independent variable and two groups, provided data followed a normal distribution and variances were equal. If variances were unequal, Welch’s two-tailed unpaired t test was applied. One-way ANOVA with Tukey post hoc correction was used for analyses with one independent variable and more than two groups, provided data followed a normal distribution. Two-way ANOVA with Sidak post hoc correction was used when two independent variables were tested. Nonparametric alternatives were used for data that did not follow a normal distribution: the Mann-Whitney U test for comparisons of two groups and the Kruskal-Wallis test with Dunn’s multiple comparisons correction for more than two groups. For cell density and regional quantifications, tests were selected based on the relevance of interactions between variables. In cases where exploring interactions was not relevant, multiple t tests or one-way ANOVA were used.

Data are presented as means ± standard deviation (SD). If multiple measurements were taken from a single individual, data are presented as means ± standard errors of the mean (SEMs). Please refer to figure legends for sample size. $p$ values < 0.05 were considered significant. Exact $P$ values are marked in the figures where space allows.

Real-time quantitative PCR (RT-qPCR).

ChP tissues were rapidly microdissected from mice, immediately frozen on dry ice, and stored at -80C. The Monarch Total RNA Miniprep Kit (New England Biolabs #T2010S) was used for total RNA isolation, eluting in 50 µL nuclease-free water. Each biological replicate consisted of a pair of lateral ventricle ChP explants or one 4$^{th}$ ventricle explant. The extracted RNA was quantified using a spectrophotometer, and 100 ng of RNA was reverse-transcribed into cDNA using the LunaScript RT Master Mix Kit (New England Biolabs #E3025L). qPCR reactions utilized specific primers and were performed in triplicate with Applied Biosystems TaqMan Gene Expression Master Mix (ThermoFisher #4369016). Probes used were Mm00487425_m1 (Fos, FAM-MGB, ThermoFisher #4331182) and Hs99999901_s1 (Eukaryotic 18s rRNA, VIC-MGB, ThermoFisher #4331182). The cycling process was carried out using the Roche LightCycler 480II. PCR cycling conditions were as follows: initial denaturation at 95°C for 10 minutes, followed by 40 cycles of 95°C for 15 seconds and 60°C for 1 minute. All reactions were performed in triplicate. Relative gene expression analysis was conducted using the 2^(-ΔΔCT) method (*33*), normalizing expression levels to 18S rRNA as the internal control.

Whole-mount clearing and light-sheet microscopy.

Tissue collection and fixation. Embryos were collected at E12.5, decapitated, and heads were immersion-fixed in 4% paraformaldehyde in PBS at 4 °C overnight. Samples were rinsed in PBS prior to SHIELD stabilization.

SHIELD stabilization (LifeCanvas, small-sample workflow). Paraformaldehyde-fixed samples were preserved with using SHIELD reagents (LifeCanvas Technologies) using the manufacturer's instructions [7]. Fresh SHIELD-OFF was prepared at 50% SHIELD-Epoxy, 25% SHIELD Buffer, and 25% water (vol/vol). Heads were incubated in SHIELD-OFF for 24 h at 4 °C with gentle shaking. SHIELD-ON was mixed with SHIELD-Epoxy at 7:1 (vol/vol); samples were incubated in this mixture for 3–6 h at 37 °C with shaking (3 h for specimens near 500 µm minimum dimension; up to 6 h for ~1.5 mm). Samples were transferred to fresh SHIELD-ON (no Epoxy) and incubated overnight at 37 °C with shaking. Tissues were stored in 1× PBS with 0.02% sodium azide at 4 °C until delipidation.

Delipidation. Three E12.5 heads were placed per 25 mL glass vial and incubated in LifeCanvas SDS-based delipidation buffer at 45 °C for 7 days on a shaking incubator set to maximal shelf motion. Buffer was refreshed according to the manufacturer's guidance. Samples were rinsed thoroughly in PBS to remove residual detergent.

Passive immunolabeling. Samples were transferred directly from delipidation buffer into blocking buffer (PBS with 0.1% Triton X-100 and 5% normal goat serum, NGS) and blocked for 2 days at 37 °C with shaking. For primary labeling, samples were moved into 10 mL of PBS + 0.1% Triton X-100 + 5% NGS containing rabbit anti-c-FOS (Synaptic Systems clone Rb108B5) at 1:500 and incubated for 5 days at 4 °C with gentle rotation. Primary wash regimen: two quick rinses in PBS + 0.1% Triton X-100, then 24 h at 4 °C with rotation in 25 mL PBS + 0.1% Triton X-100; this rinse-and-24 h step was repeated daily for 3 days. Secondary labeling was performed in 25 mL PBS + 0.1% Triton X-100 + 5% NGS with goat anti-rabbit Alexa Fluor 488 at 1:250 for 4 days at 4 °C with rotation, followed by the same wash regimen with daily solution changes for 4 days. Antibody identities and catalog numbers are listed in Materials.

Index matching. Labeled tissues were equilibrated in 50% EasyIndex and 50% water for 24 h with shaking, then in 100% EasyIndex for 24 h at room temperature with shaking. For each step, 10–20 mL of solution was used per three heads. In subsequent sessions we avoided reusing EasyIndex that had been in contact with an oil overlay, as residual oil produced refractive artifacts.

Mounting, imaging, and visualization. Samples were mounted in 100% EasyIndex in an index-matched chamber and imaged on a Zeiss Lightsheet 7 with a 5× detection objective.

Excitation was 488 nm for c-FOS with the standard emission filter set. Z-stacks spanned the full sample volume; when tiling was required, tiles were acquired with overlap and stitched in ZEN. Raw stacks were exported as 16-bit TIFFs and imported into syGlass [8] (v2.3.0; https://www.syglass.io/; RRID:SCR_017961) for volume rendering and orthoslice inspection while viewing in VR on a Meta Quest headset. A single-channel linear transfer function and identical display ranges were applied across samples. No deconvolution or selective smoothing was used.

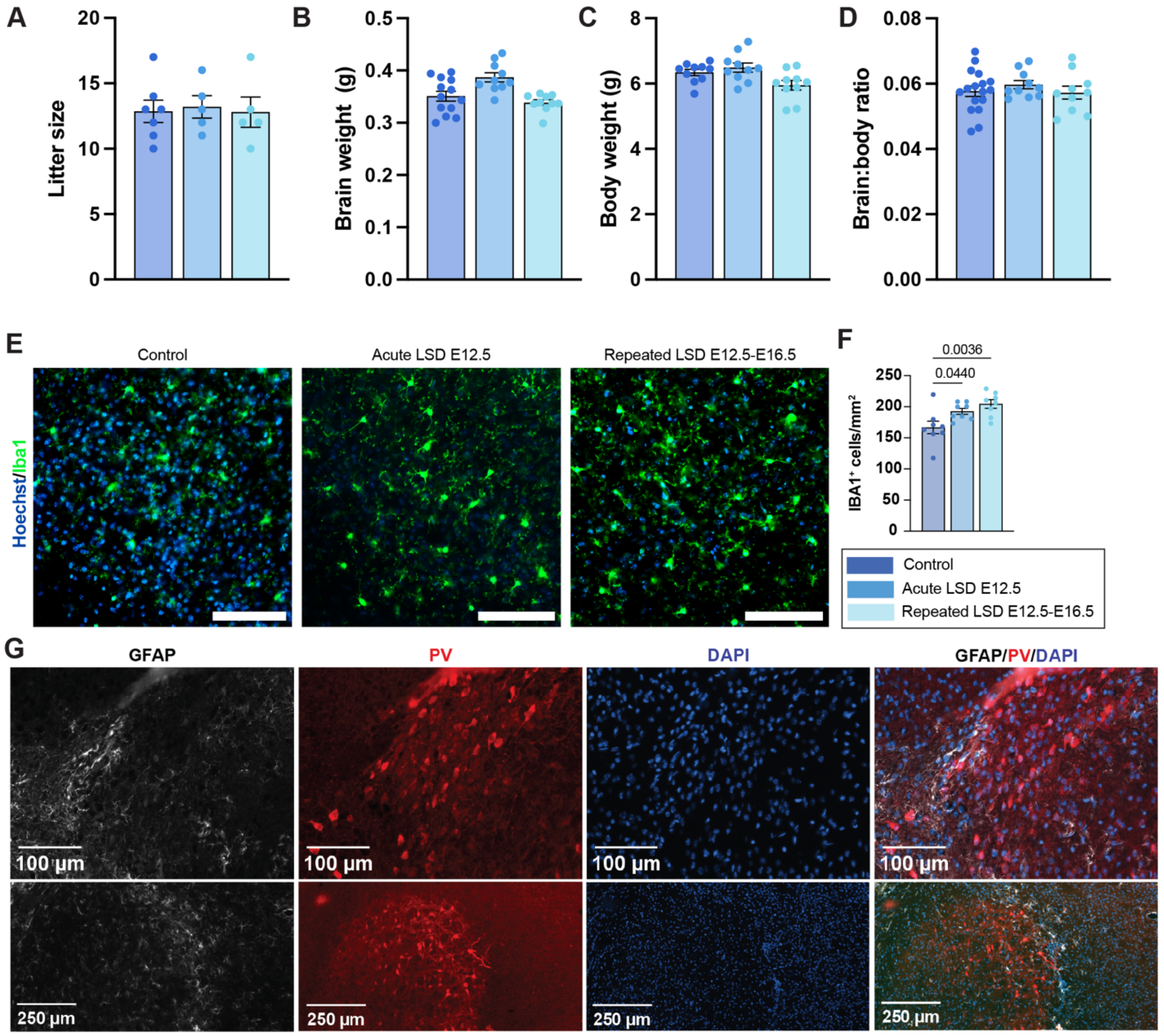

A
Litter size
B
Brain weight (g)
C
Body weight (g)
D
Brain:body ratio
E
Control
Acute LSD E12.5
Repeated LSD E12.5-E16.5
Hoechst/Iba1
F
0.0036
0.0440
IBA1+ cells/mm2
Control
Acute LSD E12.5
Repeated LSD E12.5-E16.5
G
GFAP
PV
DAPI
GFAP/PV/DAPI
100 µm
250 µm

**Supp. Fig. 1 Early postnatal growth and focal astrocyte reactivity at P8.** (**A–D**) Litter size at birth (**A**) and P8 brain weight (**B**), body weight (**C**), and brain:body ratio (**D**) for offspring from saline, acute LSD (E12.5), and repeated LSD (E12.5–E16.5) pregnancies. Points are individual pups; bars show mean ± SEM. No group differences were detected by one-way ANOVA (n.s.). **(E)** $IBA1^{+}$ microglia: representative images (scale bars, 100 µm) and (**F**) density quantification (cells $mm^{-2}$). Microglia density increases with repeated exposure and is modestly elevated after acute exposure (exact *P* values on panel). Control n = 8 from 3 litters, acute LSD n = 8 from 3 litters; repeated LSD n = 8 from 3 litters. (**G**) Coronal sections posterior to the anterior commissure stained for GFAP (astrocytes), parvalbumin (PV), and DAPI. In LSD-exposed brains, ectopic $PV^{+}$ interneuron clusters near the ventricular border are surrounded by a rim of GFAP-positive processes, consistent with local astrocyte reactivity. Scale bars: 100 µm (top), 2500 µm (bottom). Representative images are shown.

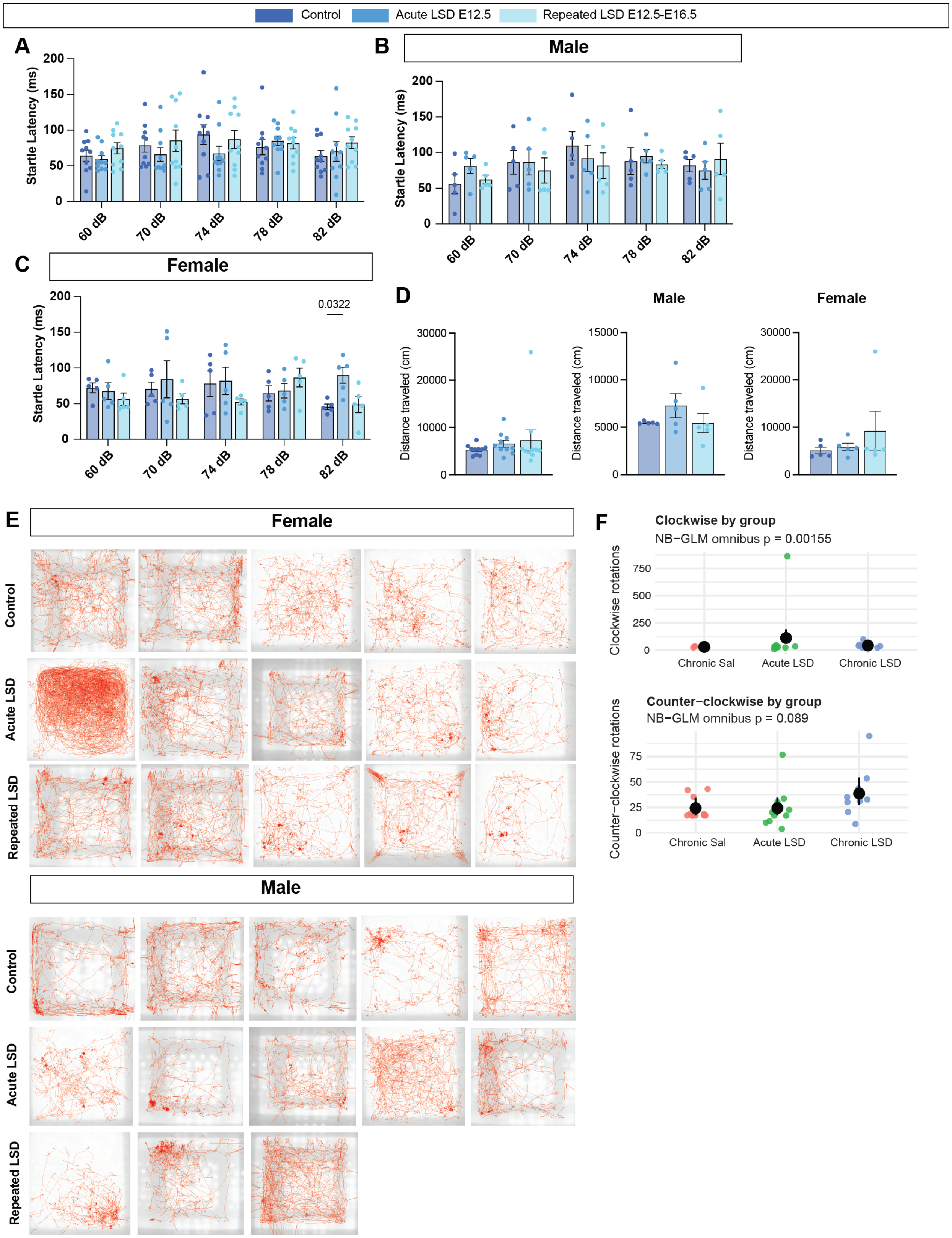
Control
Acute LSD E12.5
Repeated LSD E12.5-E16.5
A
Startle Latency (ms)
60 dB
70 dB
74 dB
78 dB
82 dB
B
Male
Startle Latency (ms)
C
Female
Startle Latency (ms)
0.0322
D
Male
Female
Distance traveled (cm)
E
Female
Control
Acute LSD
Repeated LSD
Male
Control
Acute LSD
Repeated LSD
F
Clockwise by group
NB−GLM omnibus p = 0.00155
Clockwise rotations
Chronic Sal
Acute LSD
Chronic LSD
Counter−clockwise by group
NB−GLM omnibus p = 0.089
Counter−clockwise rotations
Chronic Sal
Acute LSD
Chronic LSD

**Supp. Fig. 2 Startle latency, open-field distance, raw paths, and direction-specific rotations.** (**A–C)** Startle latency to onset (ms) across stimulus intensities (60, 70, 74, 78, 82 dB). **A**, all animals; **B**, males; **C**, females. Points are individual mice; bars are mean ± SEM. No consistent group effect is observed overall; the only reliable difference is slower responses in **acute LSD females at 82 dB** (**C**, one-way ANOVA by group at each intensity with Tukey multiple-comparison adjustment, $P = 0.0322$). **(D)** Total distance traveled (cm) in the open field shown for completeness (all animals, left; males, middle; females, right). Points are individual mice; bars are mean ± SEM. Distance is not the primary signal in this cohort. **(E)** Raw open-field path traces for all mice, separated by sex (female, top block; male, bottom block) and group (Control, Acute LSD, Repeated LSD; colors as legend). **(F)** Direction-specific rotations by group. **Top**, clockwise rotations (NB-GLM omnibus $P = 0.00155$); **bottom**, counter-clockwise rotations (NB-GLM omnibus $P = 0.089$). Points are individual mice; horizontal bars show estimated marginal means ± SE on the response scale (emmeans). Model specification and multiple-comparison procedures are detailed in Methods.

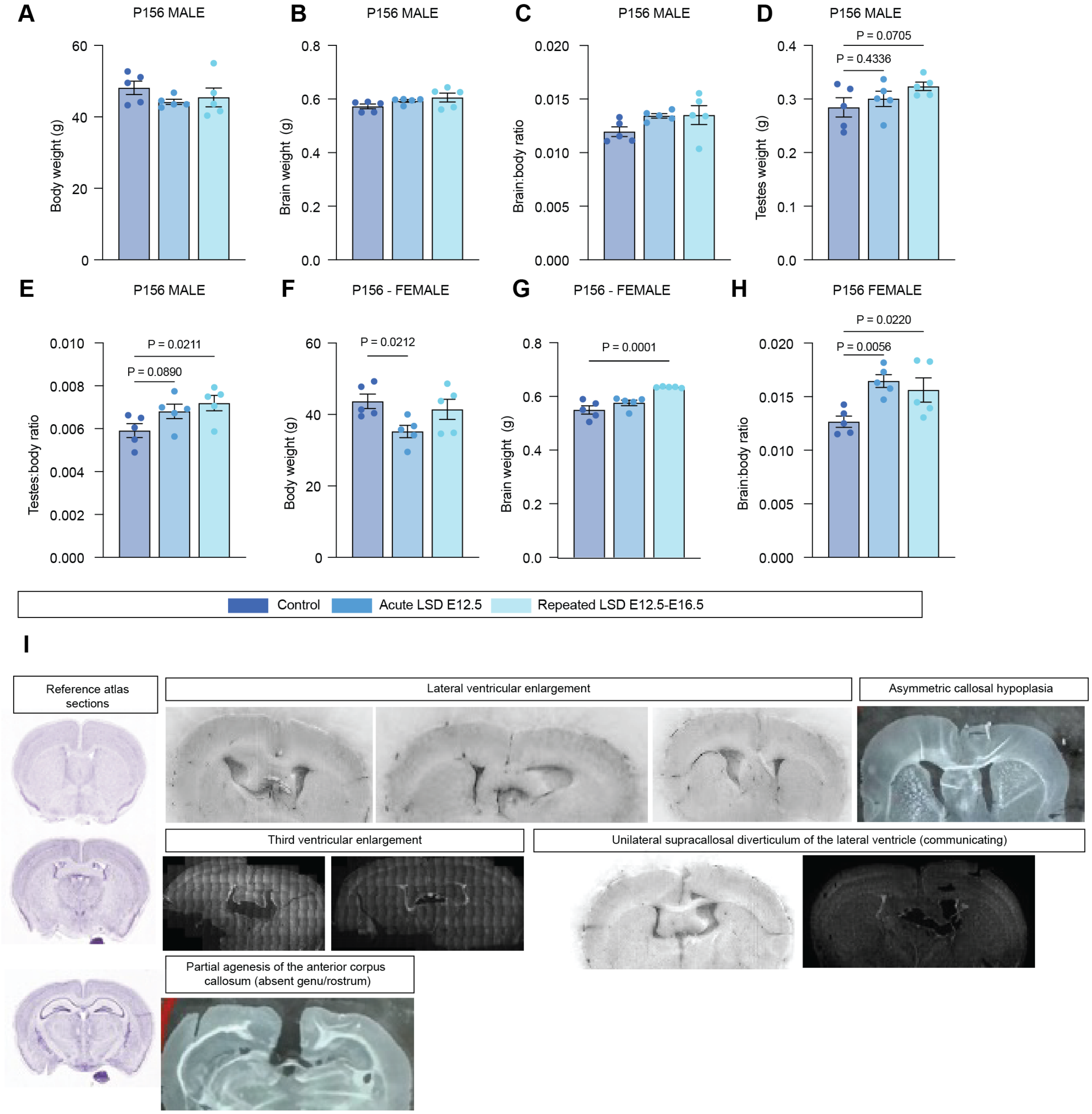
A
P156 MALE
Body weight (g)
B
P156 MALE
Brain weight (g)
C
P156 MALE
Brain:body ratio
D
P156 MALE
Testes weight (g)
P = 0.0705
P = 0.4336
E
P156 MALE
Testes:body ratio
P = 0.0211
P = 0.0890
F
P156 - FEMALE
Body weight (g)
P = 0.0212
G
P156 - FEMALE
Brain weight (g)
P = 0.0001
H
P156 FEMALE
Brain:body ratio
P = 0.0220
P = 0.0056
Control
Acute LSD E12.5
Repeated LSD E12.5-E16.5
I
Reference atlas sections
Lateral ventricular enlargement
Asymmetric callosal hypoplasia
Third ventricular enlargement
Unilateral supracallosal diverticulum of the lateral ventricle (communicating)
Partial agenesis of the anterior corpus callosum (absent genu/rostrum)

**Supp. Fig. 3 Adult (P156) anatomy after prenatal LSD.** (**A–B**) Body weight by treatment in males and females. Females: chronic E12.5–E16.5 lower than control (P=0.021); acute not significant. Males: not significant. (**C–D**) Brain weight. Females: acute E12.5 higher than control (P=0.0001); chronic not significant. Males: not significant. (**E–F**) Brain:body ratio by sex (shown as a secondary metric; not used for inference in main text). Females: higher in both LSD groups (P=0.022 and 0.0056). Males: not significant. (**G–H**) Male testes weight and testes:body ratio (acute ratio higher, P=0.021; others not significant). Points are individual animals; bars are mean ± SEM; n=5 per group per sex. Pairwise tests are Fisher's LSD following one-way ANOVAs within sex (Prism). (**I**) Midline anatomy screen at P156 (descriptive). Atlas reference levels and representative examples from LSD-exposed adults illustrate lateral ventricular enlargement with frontal-horn predominance, third-ventricle enlargement, asymmetric callosal hypoplasia, a supracallosal diverticulum continuous with the frontal horn, and partial agenesis of the anterior corpus callosum (absent genu/rostrum). Sections were cryostat-prepared for other analyses; sampling was limited and not powered for hypothesis testing.

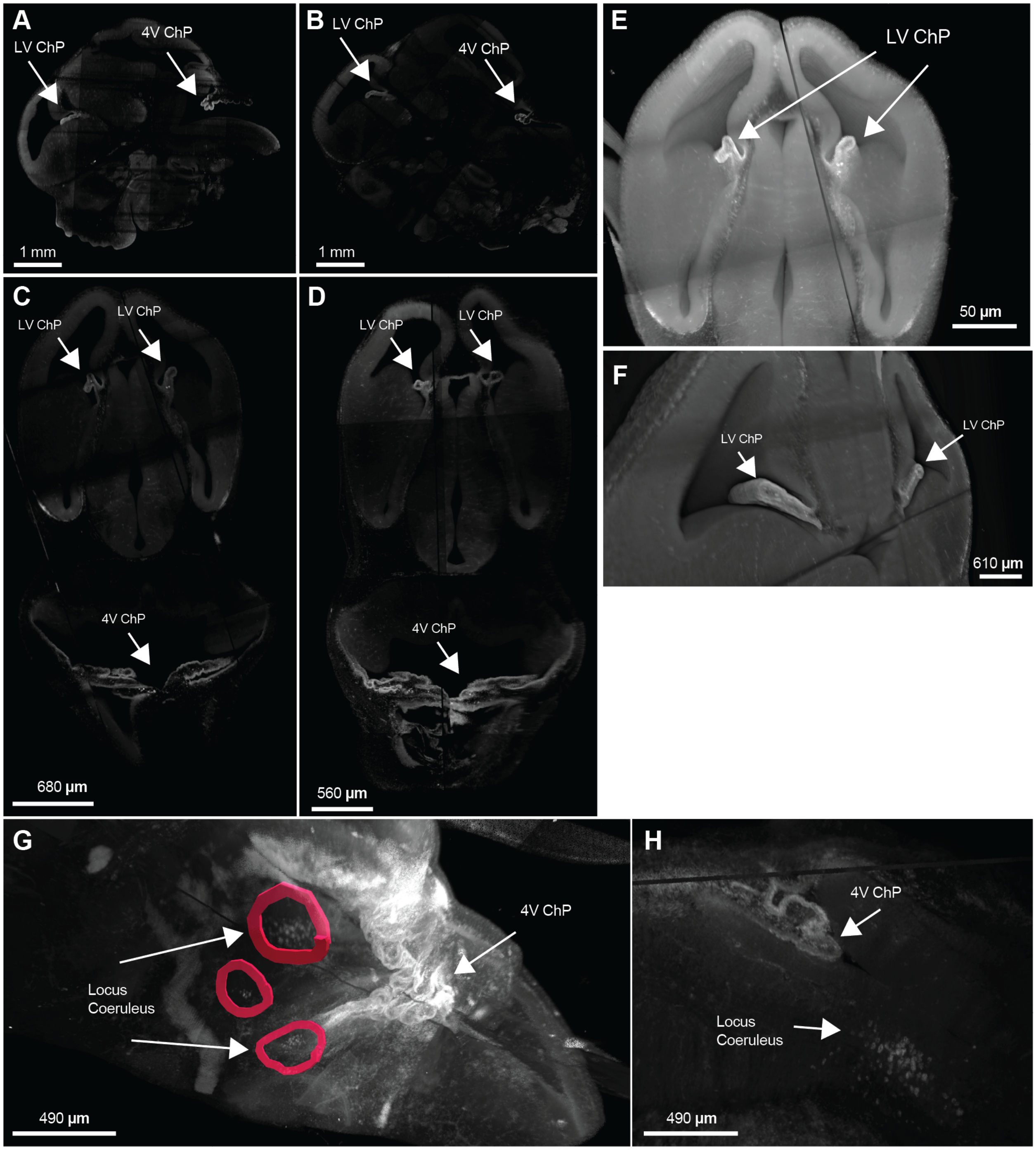
A
LV ChP
4V ChP
1 mm
B
LV ChP
4V ChP
1 mm
C
LV ChP
LV ChP
4V ChP
680 µm
D
LV ChP
LV ChP
4V ChP
560 µm
E
LV ChP
50 µm
F
LV ChP
LV ChP
610 µm
G
Locus
Coeruleus
4V ChP
490 µm
H
4V ChP
Locus
Coeruleus
490 µm

**Supp. Fig. 4 Whole-head light-sheet imaging confirms ChP activation and reveals locus coeruleus FOS signal. (A–D)** Representative cross-sectional views from optically cleared E12.5 heads 30 min after maternal LSD, stained for c-FOS. Arrows indicate **LV ChP** and **4V ChP**; robust signal is evident in **4V ChP** across views. **(E–F)** Higher-magnification coronal (**E**) and sagittal (**F**) views of LV ChP (arrows). **(G–H)** Brainstem views showing **locus coeruleus** c-FOS signal (outlined/annotated) adjacent to the 4V; shown for two independent embryos. These observations are anatomical and not quantified in the present study. Scale bars as indicated on panels. Imaging parameters and clearing protocol are in Methods. See Fig. 4L for corresponding overview images.

**Table S1**

| Analyte | Mass | Transition 1 | Transition 2 | Transition 3 |
|---|---|---|---|---|
| LSD | 324.162 | 223.00 | 208.10 | 207.10 |
| D3-LSD | 327.160 | 226.10 | 208.00 | 180.10 |

Table S1: Transitions monitored in developed LC-MS/MS method

**Table S2**

| | |
|---|---|
| Curtain Gas | 30 |
| Collision Gas | Medium |
| IonSpray Voltage | 2000 |
| Temperature | 550 |
| Ion Source Gas 1 | 30 |
| Ion Source Gas 2 | 45 |

Table S2: Source conditions optimized for analytes in developed LC-MS/MS method.

#### Supplemental References


1. Biodock, AI Software Platform. 2023.
2. Wickham H. ggplot2: Elegant Graphics for Data Analysis. 2nd ed. 2016. Cham: Springer International Publishing : Imprint: Springer; 2016.
3. Wickham H, Francois R, Henry L, Muller K, Vaughan D. dplyr: A Grammar of Data Manipulation. 2023.
4. Wickham H, Vaughan D, Girlich M. tidyr: Tidy Messy Data. 2023.
5. Courtney Y. Cortical Layer Cell Counting. 2024.
6. Byer LIJ, Xu H, Lehtinen MK. Protocol for the dissection, immunostaining, and imaging of whole-mount mouse choroid plexus. STAR Protoc. 2025;6:103627.
7. Park Y-G, Sohn CH, Chen R, McCue M, Yun DH, Drummond GT, et al. Protection of tissue physicochemical properties using polyfunctional crosslinkers. Nat Biotechnol. 2019;37:73–83.
8. Pidhorskyi S, Morehead M, Jones Q, Spirou G, Doretto G. syGlass: Interactive Exploration of Multidimensional Images Using Virtual Reality Head-mounted Displays. 2018.
